# Niche-specific contribution of the branched-chain amino acid biosynthesis protein IlvD in *Streptococcus pneumoniae* infection

**DOI:** 10.64898/2026.06.03.729751

**Authors:** Sumit Kumar, Angharad E Green, Leonie J Lorenz, Shaw Camphire, Ashima Wadhawan, Natalie O Munroe, Kyle Newbold, Graeme Ball, Emily Khlebutina, Marc S Dionne, Stephanie Lo, Daniel R Neill

**Affiliations:** Division of Molecular Microbiology, Faculty of Life Sciences, University of Dundee, Dundee, UK, DD1 5EH; Advanced Research Computing Centre (ARC), University College London, London, UK, WC1E 6EA; European Molecular Biology Laboratory-European Bioinformatics Institute, Hinxton, UK, CB10 1SD; Department of Biological Sciences, Carnegie Mellon University, 4400 Fifth Avenue, Pittsburgh, PA, USA, 15213; Department of Life Sciences and Centre for Bacterial Resistance Biology, Imperial College London, London, UK, SW7 2AZ; Dundee Imaging Facility, Faculty of Life Sciences, University of Dundee, Dundee, UK, DD1 5EH; Wellcome Sanger Institute, Wellcome Genome Campus, Hinxton, Cambridge, UK, CB10 1SA

## Abstract

Systemic pneumococcal infections are a major cause of morbidity and mortality. Pneumococci isolated from bacteraemia lack clear genetic signatures distinguishing them from isolates recovered from other disease sites. The bloodstream is an evolutionary dead end, providing no means of onward transmission, and acute bacteraemic events offer limited opportunity for within-host evolution. Nonetheless, we reasoned that it might be possible to identify genetic determinants of virulence in blood through genomic comparison of matched lung and blood isolates from individual infections. Using samples from a mouse invasive pneumonia model, we observed frequent occurrence of low population frequency mutations in *ilvD* - encoding a branched-chain amino acid (BCAA) biosynthesis protein - in bloodstream isolates. These mutations were absent from the lung-resident bacterial population. The single-nucleotide polymorphisms in *ilvD* clustered in a short stretch of nucleotides that are highly conserved across bacteria, suggesting they might disrupt protein function. Deletion of *ilvD* in pneumococcal strain D39 conferred competitive advantages in both *in vitro* environments mimicking the bloodstream and in mouse and Drosophila systemic infection models. Improved survival of *ilvD* mutants within phagocytic cells was dependent on iron availability. Conversely, disruption of *ilvD* reduced fitness in the nasopharynx, suggesting BCAA biosynthesis is required for upper airway colonisation but is a liability in the context of systemic infection. Nasopharyngeal colonisation defects could be rescued by administering intranasal BCAA to infected mice, suggesting loss of capacity for *de novo* BCAA synthesis in *ilvD* mutants accounted for reduced upper airway fitness. In support of a critical role for BCAA biosynthesis in the primary commensal lifestyle of pneumococcus, genomic analysis of clinical isolates demonstrated that *ilvD* is under purifying selection. Collectively, these findings highlight the context-specific role of IlvD and BCAA biosynthesis in infection, with a requirement for protein function in the BCAA-restricted environment of nasopharynx, but fitness benefits deriving from loss of IlvD and its associated iron-sulfur cluster, in the BCAA-rich conditions of blood.

## Introduction

*Streptococcus pneumoniae* (the pneumococcus) resides primarily as a commensal in the human nasopharynx. Most pneumococcal lineages, however, retain pathogenic potential that results predominantly from factors required by the bacterium to stimulate the inflammation necessary for onward transmission^1^. In the disease state, a changing physical and chemical environment is sensed by pneumococcus, leading to coordinated regulation of virulence factors that contribute to infection outcomes^2^.

Pneumococcus can thrive in diverse environments due to its metabolic flexibility, encoded in a wide range of transporters, enzymes and regulators. Although it primarily relies on glucose for growth, it can efficiently utilise approximately 30 different carbohydrates^3^. Distinct tissue environments exhibit unique nutrient signatures, which influence bacterial growth and virulence, ultimately shaping disease outcomes^4^.

Pneumococcus has evolved sophisticated strategies for survival in nutrient-sequestered host environments that are subject to intense immune surveillance. Expression of pneumolysin, a cytotoxic virulence factor that forms pores in cholesterol-rich host cell membranes^5^ may facilitate release of host-derived nutrients, enabling bacterial scavenging. In the nasopharynx, pneumococcus acquires sugars from glycosylated mucins through glycoside hydrolases and specialised sugar transport systems. Allelic variants of metabolic genes and in the genetic architecture of promoter regions further contribute to lineage-specific success in different host nutritional environments ^2,6,7^. Such variations fine-tune transcriptional and translational control of sugar, metal, and amino acid transport systems, allowing the bacterium to acquire fundamental biochemical building blocks whilst reducing the requirement for *de novo* synthesis.

Iron acquisition is indispensable for pneumococcal survival; however, unlike many respiratory pathogens, it lacks known siderophores. Instead, pneumococcus scavenges iron via surface proteins such as PspA, which binds to the iron-sequestering host glycoprotein lactoferrin^8^, as well as via the actions of pneumolysin on host cells, the uptake of ferric iron by PiaABC and PiuABC transport systems and uptake of heme-bound iron via PitABC^9,10^. Pneumococcus requires iron for DNA synthesis and repair and as a cofactor for enzyme function in proteins containing iron-sulfur clusters. Pneumococcus has relatively few Fe-S cluster proteins, likely due to the endogenous production of hydrogen peroxide by pneumococcal SpxB and the consequent risk of catalysing Fenton chemistry. However, several pneumococcal Fe-S cluster-containing proteins play key roles in central and anaerobic metabolism^11^.

The branched-chain amino acids (BCAAs), isoleucine, leucine, and valine, are also essential for pneumococcal growth and can be scavenged from the host via transporters, including LivJHMGF and BrnQ ^12^. Pneumococcus contains all the biosynthetic machinery required for *de novo* BCAA synthesis, including the dihydroxy-acid dehydratase IlvD, which contains a 4Fe-4S cluster^13,14^. However, *in vitro* pneumococci behave as apparent BCAA auxotrophs^15^. This dichotomy may reflect challenges in replicating the nutritional conditions required to de-repress the BCAA biosynthetic gene cluster *in vitro*. Within-host, BCAA availability varies significantly across tissue sites. BCAAs are scarce in the primary nasopharyngeal niche, are found at modestly higher concentrations in lungs^4^ and are abundant in blood^16^, suggesting tight regulation of transporters and biosynthetic pathways may be necessary.

In this study, we identified the regular occurrence of low-frequency non-synonymous *ilvD* mutations in populations of pneumococci isolated from blood in a mouse invasive pneumonia model. Mutations in *ilvD* were not observed in matched lung pneumococcal populations from the same mice. When identified SNPs were reproduced in the laboratory *S. pneumoniae* strain D39, the resulting mutants partially phenocopied an *ilvD* deletion strain. To determine whether the absence of IlvD confers a survival advantage in the bloodstream, we assessed pneumococcal growth in blood, survival in phagocytic cells and virulence in a mouse bacteraemia model. Mutants in *ilvD* showed enhanced virulence in blood and prolonged survival within phagocytic cells under iron-limited, but not iron-replete conditions. By contrast, loss of IlvD attenuated both colonisation and virulence potential in the BCAA-limited environment of the airways. Notably, this phenotype was reversible with exogenous BCAA supplementation in a mouse nasopharyngeal carriage model. We hypothesise that reduced IlvD expression in BCAA-replete environments, such as blood, is beneficial via reduced energy demands and by enabling prioritisation of iron for Fe-S cluster-containing proteins required for anaerobic growth or oxidative stress resistance. Observed fitness defects for *ilvD* mutants in the airways suggest that *de novo* synthesis of BCAAs is required in these nutrient-scarce environments and that previous observations of BCAA auxotrophy in pneumococci are likely *in vitro* artefacts.

## Results

### Paired analysis of lung and blood bacterial populations reveals frequent mutation of *ilvD* in blood-borne pneumococci

In a previous pneumococcal experimental evolution study, we used a mouse model of pneumonia to explore adaptation of our in-house D39 laboratory reference strain (D39N) to the environment of the lower respiratory tract^17^. In 46 of 50 infected mice, pneumonia progressed naturally to bacteraemia. In these animals, pneumococci were recovered separately from lungs and blood upon euthanasia. Short-read sequencing was performed on both populations, with mutations identified using the Breseq pipeline, run in population mode^18^. This enabled paired analysis of lung and blood pneumococcal populations for identification of (1) variants or sub-populations arising uniquely in blood, (2) clones present in lung but lost upon transition to blood via bottlenecks or selection, and (3) clones appearing first in lung but expanding upon transition into blood. Most diversity present in lung populations - relative to the ancestor D39N strain from which they were derived – was maintained in the blood bacterial population, suggesting either that minimal bottlenecks are associated with the transition to bacteraemia in this model, or else that, once the lungs are damaged, repeated seeding events into blood can occur. Of the 143 mutations identified in lung populations, 29 (20%) were not present in the matched blood populations (Figure 1A). None of those 29 mutations were fixed in the lung populations, instead being present in sub-clones at frequencies between 5% (the limit of detection at the sequencing depth achieved) and 54%. Of 138 mutations identified in blood pneumococcal populations, 24 were not found in lung populations, and these ranged from 5% to 31% population frequency (Figure 1B, Supplementary Figure 1, Supplementary Dataset 1). These mutations either arose in blood or were present at undetectable frequency amongst the lung population and expanded following seeding into the bloodstream. Amongst the 24 such mutations identified were point mutations in genes encoding nucleotide transporters, regulators of lipid and amino acid metabolism, and proteins involved in cell wall biogenesis. Of note, five unique, but low frequency (<10%) non-synonymous mutations were identified in *ilvD*, each occurring in separate blood populations, indicating they arose or were selected independently and suggesting they may confer an advantage during bacteraemia. The mutations spanned a short stretch of the coding sequence that is highly conserved in *ilvD* across bacterial species^19,20^ (Figure 1C), and resulted in sub-clones with S273C, T274P, N275I, T277N and L278I amino acid substitutions within IlvD. Given the strong nucleotide sequence conservation in this region of the gene, we hypothesised that the mutations might result in loss of IlvD function.

**Figure 1.**
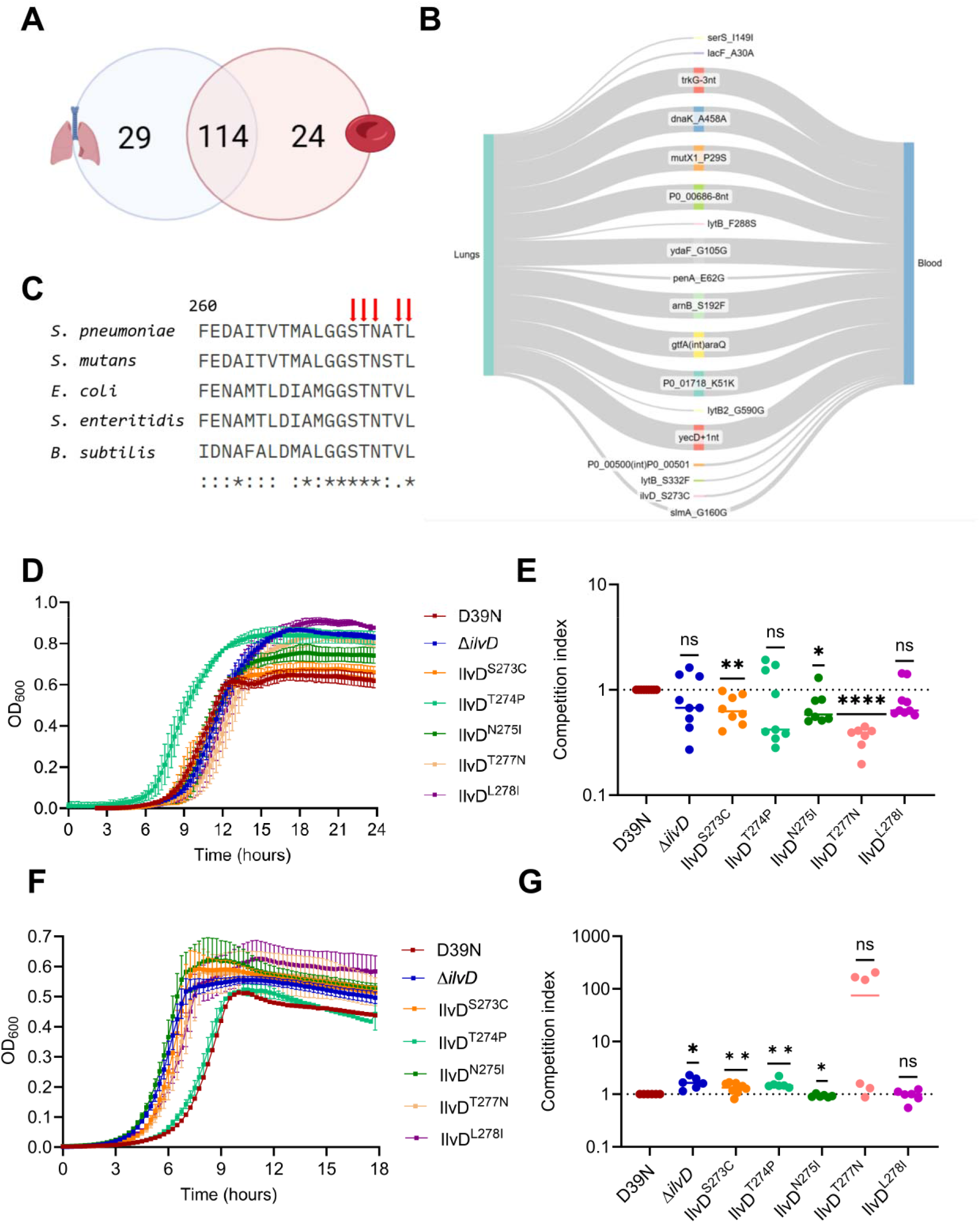
Non-synonymous mutations in *ilvD* in pneumococci isolated from bacteraemia. **A.** Mutations in lung and blood pneumococcal populations collected from a mouse invasive pneumonia model. All mutations present at greater than 5% population frequency are included. Mutations were called against the parental D39N strain and analysis is from 50 matched lung and blood populations. **B.** Sankey plot showing the fate of pneumococcal genotypes identified in lungs during the progression to bacteraemia. Lines originating in lungs and terminating at the midpoint represent diversity not carried over to blood. Lines arising at the midpoint represent diversity present in blood but not identified in lungs. The thickness of the line represents the relative frequency of each mutation. The plots shown is a representative example of matched lung and blood bacterial populations from one of ten experimental-evolution lineages, originally described in Green *et al.*^17^. Plots generated using SankeyMATIC. **C.** Protein sequence alignment of a region of IlvD, beginning from amino acid 260, in which non-synonymous mutations were identified. Arrows represent sites of identified mutations. Asterisks represent perfectly conserved residues across the species represented. Alignment generated with Clustal Omega. **D.** Growth in BCAA-limited chemically defined media (CDM) (5 µg/mL of each BCAA). **E.** The streptomycin-resistant D39N (D39N^SmR^) was competed against *ilvD* mutant strains in a 1:1 ratio in BCAA-restricted CDM. Competition indices were adjusted to account for the fitness disparity between D39N and D39^SmR^. Data was analysed by Wilcoxon t test vs a neutral competition index of 1. **F.** Growth in BHI. **G.** The streptomycin-resistant D39N (D39N^SmR^) was competed against *ilvD* mutant strains in a 1:1 ratio in BHI. Competition indices were calculated and adjusted by a factor of 0.88 to account for the fitness ratio of D39N to D39N^SmR^. Data analysis is by Wilcoxon t test vs a neutral competition index of 1. *\*p*=0.05, *\*\*p*=0.001, *\*\*\*p*=0.0001, *\*\*\*\*p*<0.0001, *ns*=non-significant.

### Mutations in IlvD confer increased fitness in serum and other nutrient replete environments

To determine whether the identified *ilvD* mutations resulted in loss of protein function, we reproduced each in SNP in isolation on a D39N background and additionally generated an in-frame deletion of the full length *ilvD* gene. We first cultured *ilvD* mutants and wild type D39N in BCAA-free chemically defined media (CDM), but consistent with previous reports of BCAA auxotrophy in pneumococci^15^, we observed no growth for any strain. To instead simulate conditions of BCAA-scarcity, we performed growth assays in CDM with BCAAs diluted 500-fold (to 5 µg/L), observing no significant difference in growth rates between the strains (Figure 1D). When *ilvD* mutants were individually competed in BCAA-restricted CDM against a D39N strain carrying a point mutation that confers streptomycin resistant (D39N^SmR^, carrying mutation RpsL^K56T^), competitive fitness was found to be modestly impaired for three of the *ilvD* SNP strains (Figure 1E). By contrast, in both single-strain growth and competition assays in nutrient broth (brain heart infusion [BHI]), *ilvD* mutants showed a clear fitness advantage relative to D39N. All strains except the IlvD^T274P^ mutant showed a shortened lag time in BHI, relative to D39N (Figure 1F) and the Δ*ilvD* strain and three of five SNP mutants outcompeted the wild type in competition assays in nutrient broth (Figure 1G). These findings demonstrate that the relative cost or benefit of *ilvD* mutations depends upon environmental conditions and BCAA availability.

### Loss of IlvD enhances virulence in blood and survival of phagocytosis

As *ilvD* mutations were identified exclusively in the blood-borne pneumococcal population, we reasoned that they must provide fitness benefits of relevance to systemic infection. To investigate this, we first assessed the competitive fitness of *ilvD* SNP and deletion strains in human serum, a BCAA-replete environment, with concentrations typically ranging from ∼200–800 µM^16^, likely reducing the need for endogenous synthesis. When *ilvD* mutants were competed against D39N^SmR^ in serum, we observed a competition index >1 in all tested strains, with significantly increased competitive fitness recorded for Δ*ilvD,* IlvD^S273C^ and IlvD^T274P^ (Figure 2A).

**Figure 2.**
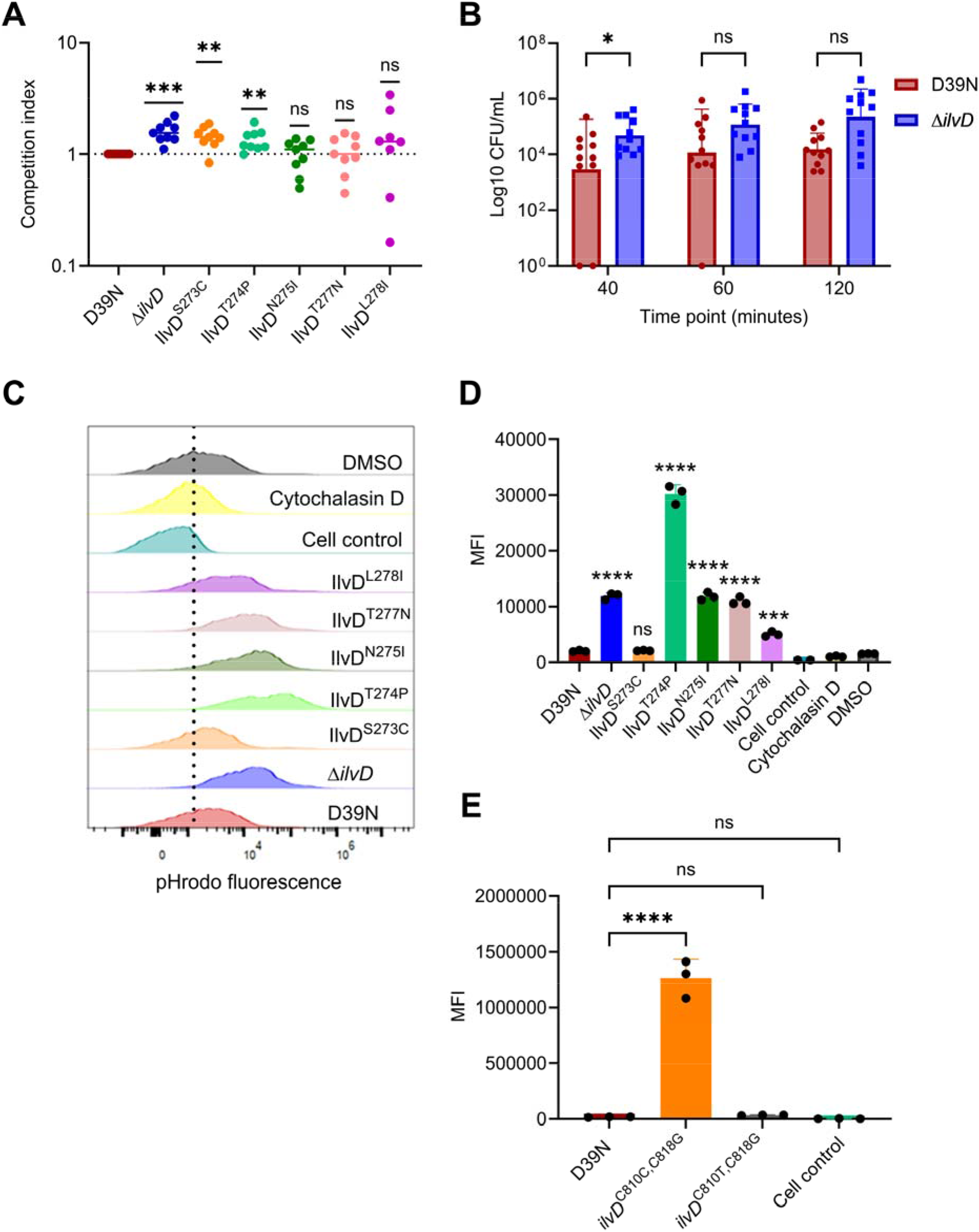
Mutations in *ilvD* are associated with increased fitness in blood. **A.** D39N^SmR^ was competed against *ilvD* mutant strains in a 1:1 ratio in human serum. Competition indices were adjusted to account for the fitness ratio of D39N to D39N^SmR^. Data analysis is by Wilcoxon t test vs a neutral competition index of 1. **B.** Phagocytosis assay was performed using PMA-activated THP-1 cells. THP-1 surface associated and intracellular pneumococci were recovered from washed and lysed cells at 40, 60 and 120 minute time points. Data analysis is by two-way ANOVA with Sidak’s multiple comparison test. **C.** Bacteria were labelled using the pH-sensitive dye pHrodo before infecting THP-1 cells. After 60 minutes of incubation, infected cells were analysed by flow cytometry. **D.** Median fluorescence intensity (MFI) obtained from THP-1 cells infected with pHrodo-labelled bacteria. Data analysis is by one-way ANOVA. **E.** Repeat pHrodo assay using two clones of the IlvD^S273C^ mutant. One carries only the mutation at the 818^th^ nucleotide that was identified in the blood-borne pneumococcal population. The other carries an additional nucleotide change introduced during or subsequent to cloning. *\*p*=0.05, *\*\*p*=0.001, *\*\*\*p*=0.0001, *\*\*\*\*p*<0.0001, *ns*=non-significant.

A key host immune defence against systemic infection is mediated through the action of phagocytic cells. Previous studies using a *Streptococcus mutans* strain with disruptive mutation of *ilvE* reported enhanced intracellular survival within pHSC-2 cells, relative to wild type^21^. As IlvE functions in the same biosynthetic pathway as IlvD, catalysing the final step in BCAA synthesis, we assessed uptake and survival of the Δ*ilvD* strain in phagocytic THP-1 cells. We recovered increased numbers of viable pneumococci from THP-1 cells infected with the Δ*ilvD* strain, relative to D39N, at all tested timepoints (Figure 2B). To determine whether the *ilvD* SNP mutants phenocopied the deletion strain, we employed the acid-sensitive dye pHrodo, which fluoresces specifically in acidic environments, such as the phagosome. THP-1 cells were infected with pHrodo-labelled bacteria and fluorescence assessed after 60 minutes. Cells infected with the Δ*ilvD* strain and 4 of 5 *ilvD* SNP mutants exhibited increased fluorescence, relative to cells infected with D39N, indicating improved uptake or prolonged survival within THP-1 cells (Figure 2C, D). Cells infected with the IlvD^S273C^ strain displayed a fluorescence profile similar to those infected with D39N. To investigate this, we re-sequenced the *ilvD* gene from two separate clones of this mutant. This revealed that the clone used in our phagocytosis assays had acquired a second – apparently compensatory – mutation in *ilvD*, at nucleotide position 810 (C810T). We repeated the pHrodo assay with the second IlvD^S273C^ clone, lacking the C810T mutation and observed a fluorescence profile matching those of the *ilvD* deletion strain and other SNP mutants (Figure 2E).

### Loss of IlvD enhances virulence in a mouse bacteraemia model but not in pneumonia

As *ilvD* mutants showed evidence of enhanced fitness in serum and survival of phagocytosis, we reasoned they might display enhanced virulence in the bloodstream. We performed intravenous infections with D39N or Δ*ilvD* and monitored disease progression over 72 hours (Figure 3A). Mice infected with the Δ*ilvD* strain developed disease symptoms earlier and succumbed faster than D39N-infected mice, although survival curves were not significantly different in Kaplan-Meier analysis. Increased counts of viable bacteria from blood were recorded for the Δ*ilvD* infections, relative to D39N infections, at all tested timepoints (Figure 3B). The heart has been suggested as a protected environment within which pneumococci form microlesions during systemic infection^22^. Comparison of pneumococcal burden in cardiac tissue at 24 hours post-infection revealed increased numbers of heart-resident bacteria in Δ*ilvD*-infected mice (Figure 3C). Collectively, these findings suggest a more rapid *in vivo* expansion of the Δ*ilvD* strain during bacteraemia, relative to D39N, mirroring the *in vitro* results in BCAA-replete media and those from competition assays in human serum.

**Figure 3.**
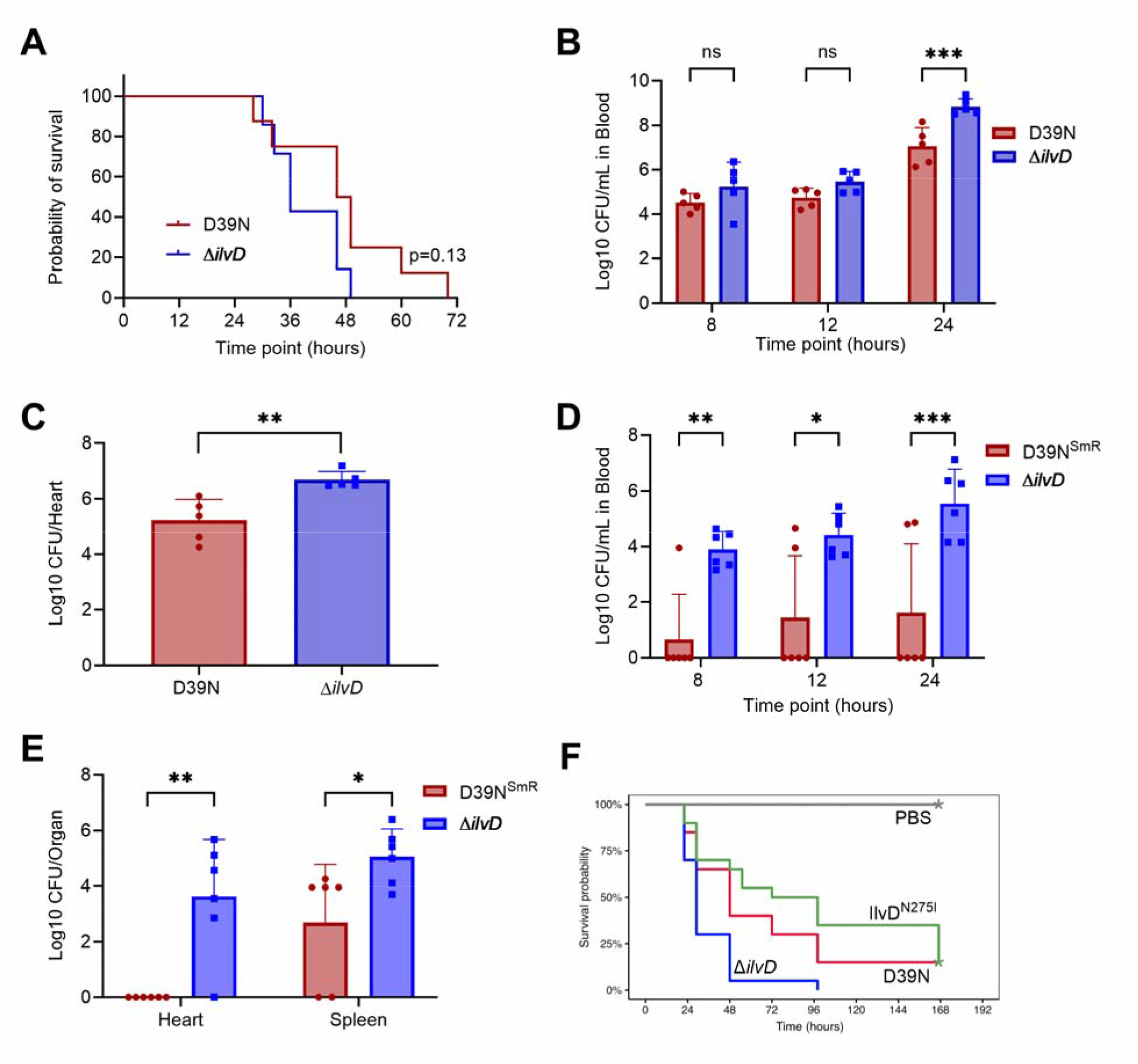
Loss of IlvD is associated with increased virulence in a mouse bacteraemia model. **A.** Survival in a mouse intravenous infection model, n=8 mice per group. P value determined by Kaplan-Meier analysis. **B.** Blood colony forming units (CFU) over a 24 hour timecourse, n=5 mice per group. **C.** Bacterial recovery from heart tissue at 24 hours post infection. **D.** Mice were intravenously administered a mix of D39N^SmR^ and D39NΔ*ilvD* in equal proportions and the CFU in blood for each strain was determined at 8, 12 and 24 hours post infection by parallel plating of blood samples on streptomycin and non-selective agar. **E.** Colony forming units of D39N^SmR^ and D39NΔ*ilvD* recovered from heart and spleen tissue at 24 hours post-infection in the mixed-strain intravenous infection model. **F.** Survival of *Drosophila melanogaster* in an acute infection model. Data are a composite of three independent experiments. Analysis in **B., D.** and **E.** is by two-way ANOVA with Sidak’s multiple comparison correction. Analysis in **C.** is by unpaired t-test. *\*p*=0.05, *\*\*p*=0.001, *\*\*\*p*=0.0001, *\*\*\*\*p*<0.0001, *ns*=non-significant.

To determine whether the human serum competition results could be reproduced *in vivo*, we performed a mixed infection experiment in CD1 mice using both D39N^SmR^ and the Δ*ilvD* strain. Analysis of bacterial loads in blood at 8, 12, and 24 hours post-infection demonstrated that the *ilvD* deletion strain outcompeted D39N^SmR^ (Figure 3D). No viable D39N^SmR^ bacteria were recovered from blood for four of the six infected mice. Analysis of heart and spleen – two environments that act as protected reservoirs of pneumococci during systemic infection ^22,23^ – at 24 hours post-infection revealed higher bacterial loads for the Δ*ilvD* strain. Notably, however, D39N^SmR^ cells were recovered from the spleens of four of six infected animals, supporting previous observations that viable pneumococci can be maintained within the spleen even when the bloodstream has been sterilised by the actions of circulating phagocytic cells^23^ (Figure 3E). Finally, to test the impact of *ilvD* mutations on virulence in an infection model in which phagocytes represent the primary cellular immune defence, we compared D39N, the *ilvD*-deletion strain and the IlvD^N275I^ mutant in a *Drosophila melanogaster* infection model^24^. Under these conditions, the Δ*ilvD* strain, but not the SNP mutant, showed enhanced virulence relative to wild type D39N (Figure 3F).

As the *ilvD* mutations we had identified in blood-borne pneumococci were not observed in matched lung bacterial populations, we sought to determine whether the increased *in vivo* fitness of the Δ*ilvD* strain would be retained in a pneumonia model that progresses naturally to bacteraemia. In this model, strains must first establish an infection within the more BCAA-restricted environment of lungs^4^ before breaching epithelial barriers and establishing systemic disease. The Δ*ilvD* strain retained capacity to cause invasive pneumonia but did not show the fitness advantage, relative to D39N, that had been observed in the intravenous infection model (Figure 4). No bacteria were identified in the blood of any mice at 8 hours post-infection in this model, whilst bacterial loads in blood at both 12 and 24 hours post-infection were comparable for Δ*ilvD*-and D39N-infected mice (Figure 4A). There was a non-significant trend towards reduced viable bacterial numbers in heart and lungs of Δ*ilvD*-infected mice (Figure 4B) and analysis of bacterial microlesions in tissue histology sections revealed a higher number of such plaques within the hearts of D39N-infected mice, relative to those infected with the Δ*ilvD* strain (Figure 4C, E).

**Figure 4.**
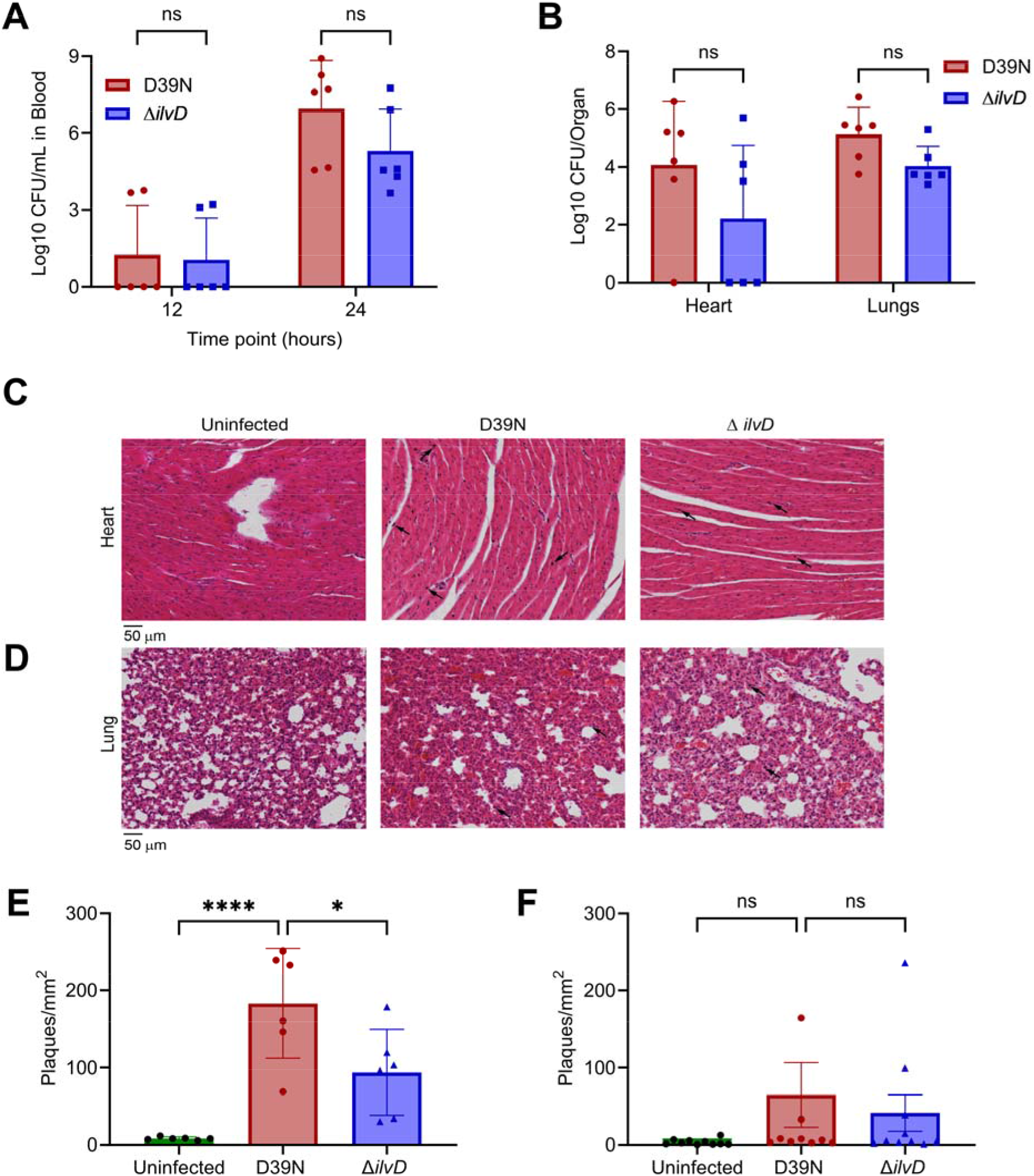
IlvD is not essential for establishment of pneumonia but may contribute to fitness in lungs. **A.** Blood colony forming units (CFU) at 12 and 24 hours post intranasal infection with a pneumonia-inducing dose of D39N or D39NΔ*ilvD*, n=6 mice per group. **B.** Bacterial CFU in heart and lung tissue at 24 hours post-infection. **C.** Hematoxylin and eosin (H&E)-stained heart sections from D39N-and D39NΔ*ilvD*-infected mice and uninfected controls, showing presence of pneumococcal plaques in infected tissue (black arrows). **D.** H&E-stained lung sections, with pneumococcal plaques shown. **E.** Quantification of pneumococcal plaques from heart histology. **F**. Quantification of plaques from lung histology. Analysis is by two-way (**A., B.**) or one-way (**E., F.**) ANOVA, with multiple comparison correction. *\*p*=0.05, *\*\*\*\*p*<0.0001, *ns*=non-significant.

### IlvD is required for prolonged nasopharyngeal carriage

IlvD is highly conserved amongst bacteria, suggesting a pivotal role in microbial physiology^19^. However, pneumococci are auxotrophic for BCAAs *in vitro* and the *ilvD* SNPs we identified in bloodstream populations appeared to be loss of function mutations. We therefore investigated whether IlvD was dispensable for pneumococci within their natural nasopharyngeal niche. A mixture of D39N^SmR^ and Δ*ilvD* were instilled into the nares of mice in equal proportions, at a density that establishes nasopharyngeal carriage with only a small proportion of mice (3-5%) going on to develop pneumonia or invasive disease. Bacterial densities in nasopharynx were monitored over 14 days (Figure 5A). The bacterial burden in nasopharynx at day 1 post-infection comprised equal proportions of both strains, but at later timepoints the D39N^SmR^ strain made up an increasing proportion of the overall bacterial load. Half of the infected mice had cleared the Δ*ilvD* strain by day 7 and no viable Δ*ilvD* bacteria were recovered from nasopharynx at day 14 post-infection.

**Figure 5.**
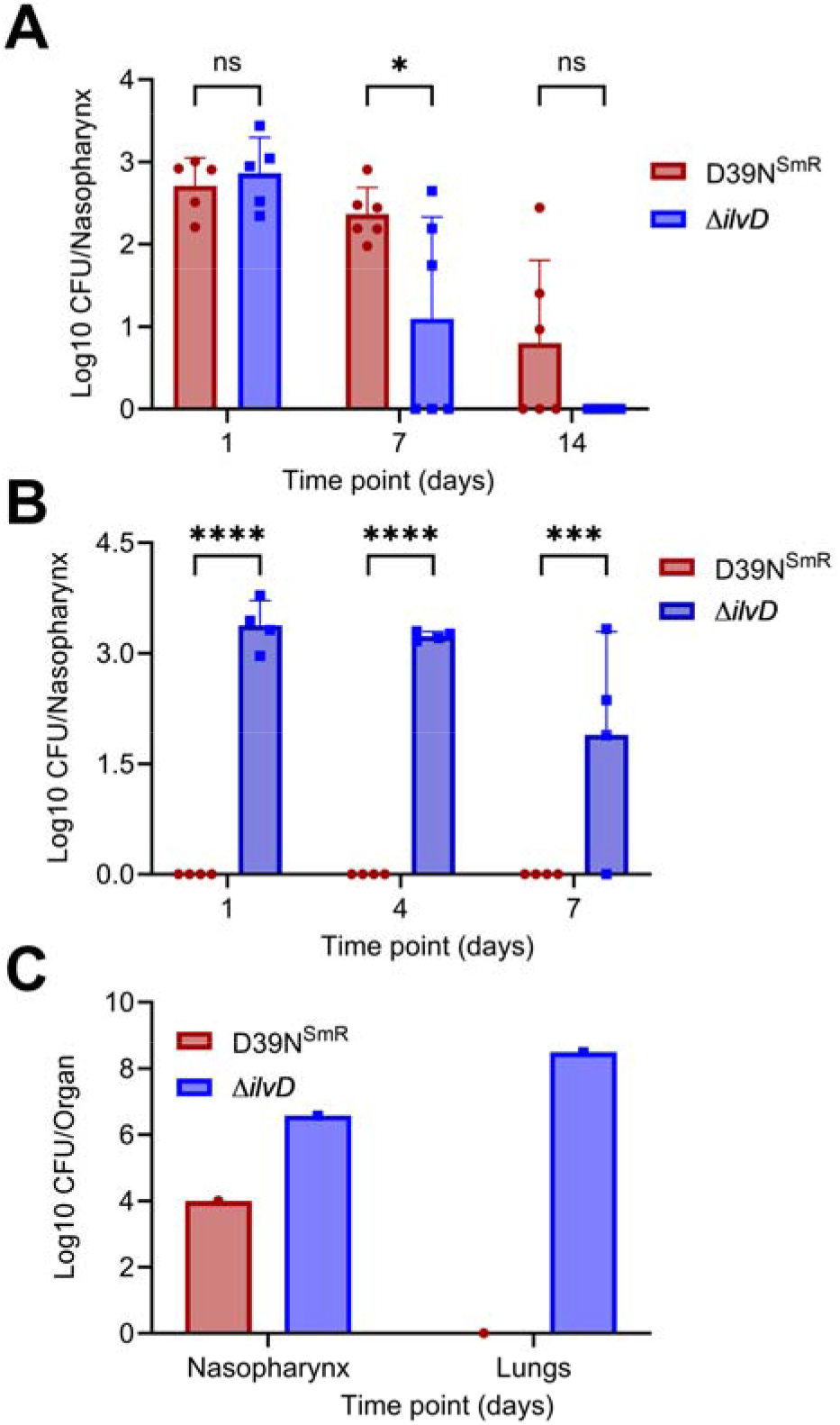
IlvD is required for long-term nasopharyngeal carriage. **A.** Bacterial colony forming units (CFU) in nasopharynx of mice intranasally infected with a low dose (1×10^5^ CFU) of D39N^SmR^ or D39NΔ*ilvD*. n=5 mice per group. **B.** Bacterial CFU in nasopharynx of mice intranasally infected with a mixed dose of D39N^SmR^ and D39NΔ*ilvD* in equal proportions, n=4 mice per group. CFU determined by parallel plating of tissue homogenates on streptomycin and non-selective agar. **C.** Bacterial CFU in nasopharynx and lungs of a mouse from the experiment described in **B.** that developed symptomatic disease at day three post-infection and was culled due to ill health. Analysis is by two-way ANOVA, with Sidaks multiple comparison correction. *\*p*=0.05, \*\*\**p<*0.001*, ****p*<0.0001, *ns*=non-significant.

We reasoned that the more rapid clearance of the *ilvD* deletion strain from the nasopharynx might reflect a failure to maintain adequate BCAA production or uptake in an environment of limited BCAA availability. We repeated the co-infection experiment, this time administering the infection dose in a 10 mg/ml BCAA solution. Further intranasal administration of BCAA solution was performed on days 2, 4 and 6 post-infection. Exogenous BCAA supplementation reversed the phenotypes observed in the original colonisation experiment, with the Δ*ilvD* strain rapidly outcompeting D39^SmR^ (Figure 5B). In this experiment, one mouse developed pneumonia at day 3 post-infection and was euthanised. We recovered both Δ*ilvD* and D39N^SmR^ from the nasopharynx of this animal (Figure 5C), confirming that the D39N^SmR^ strain was not completely cleared, although it remained below the limit of detection in the nasopharynx of all mice that did not develop systemic disease (Figure 5B). Interestingly, only the Δ*ilvD* strain was recovered from the lungs of the mouse that developed pneumonia (Figure 5C), suggesting the deletion strain was the causative agent of disease.

### Genomic analysis of clinical isolates demonstrates *ilvD* is under purifying selection

We profiled disruptive and non-synonymous mutations in *ilvD* across a collection of 24,455 pneumococcal genomes, comprising isolates from 59 countries^25^. We identified non-synonymous mutations in 9,744 genomes (40%) (Supplementary Figure 2), frameshifts introducing premature stop codons in three genomes, and three genomes containing nonsense mutations (Supplementary Table 1). The low frequency SNPs identified in blood-borne bacterial populations from the mouse studies were not identified as fixed mutations in any clinical isolates.

We next tested whether any non-synonymous mutations were associated with invasive pneumococcal disease (IPD) (n = 6133) or carriage (n = 2835). Five mutations were significantly associated with IPD isolates (Table 1). S38P and V428I were the most common non-synonymous mutations and were observed in multiple lineages, whilst V75I was associated predominantly with Global Pneumococcal Sequencing Cluster (GPSC)26-serotype 12F - a lineage with a high propensity to cause IPD^26^.

**Table 1.**
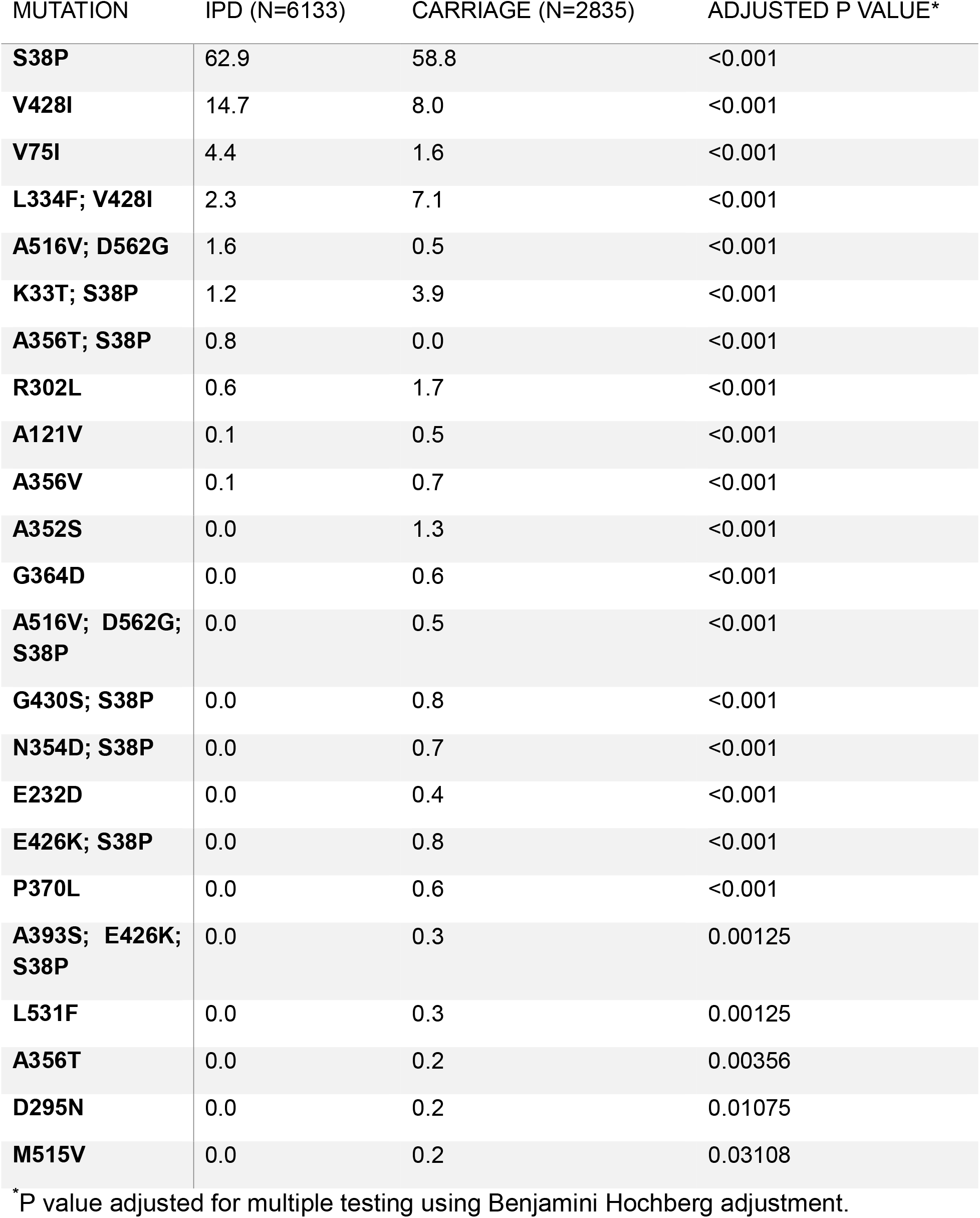
Mutations in *ilvD* that are significantly enriched amongst IPD isolates.

To explore the evolutionary pressures acting on *ilvD*, we assessed the ratio of non-synonymous to synonymous mutations at each amino acid position across the length of the protein, amongst clinical isolates. Median dN/dS values <1 for 510 of the 567 amino acid positions indicated that *ilvD* is predominantly under purifying selection. For the remaining 57 positions, although median dN/dS values were greater than one, the 90% credible intervals included one (Supplementary Figure 3, Supplementary Dataset 2). These findings align with our observation that *IlvD* is an important determinant of pneumococcal fitness in its primary nasopharyngeal niche. Identified fixed disruptive mutations amongst IPD isolates may be the result of rapid selection of previously rare mutations during bacteraemia.

### Loss of IlvD confers increased fitness in iron-limited environments

The mechanistic explanation for the niche-specific fitness benefit conferred by loss of IlvD remains somewhat elusive. The SNPs identified in the experimental evolution study partially phenocopied the *ilvD* deletion strain, but differences in strain fitness were apparent in some assays, leaving the possibility that some residual protein function remains in strains carrying those mutations. Structural predictions for IlvD show three predicted binding sites for magnesium ions, one of which sits within a protein pocket between aspartic acid 126 and glutamic acid 448. We identified a SNP at asparagine 275, which forms two hydrogen bonds with the glutamic acid at position 448 (Supplementary Figure 4). Mutations at this position may destabilise the binding of metal cofactors, compromising protein function.

To understand the fitness impacts of loss of *ilvD* we looked to the pneumococcal capsule, given its role as a key anti-phagocytic defence. Increased capsule thickness reduces uptake of pneumococcus by phagocytic cells but might also provide resistance to intracellular killing for those cells that are successfully engulfed. However, we observed no significant difference in capsule thickness between wild type and Δ*ilvD* pneumococci (Supplementary Figure 5).

We reasoned that improved competitive fitness in blood and increased survival within phagocytic cells may result, in part, from increased growth rates of the Δ*ilvD* strain in BCAA-replete environments (Figure 1F,G, Figure 2A). In support of this, the BCAA transporter gene *livJ* showed evidence of increased expression in the Δ*ilvD* strain, relative to wildtype, based on RT-qPCR analysis performed on RNA extracted during growth in whole human blood (Supplementary Figure 6). A second BCAA transport gene, *brnQ,* showed a non-significant trend towards increased expression in the *ilvD* deletion strain. This may suggest a switch away from *de novo* synthesis and towards uptake of exogenous BCAAs for the Δ*ilvD* strain during growth in blood. Another contributor to observed phenotypes may be altered activity of the major transcriptional regulator CodY that controls gene expression as a function of BCAA bioavailability^27^. However, whilst expression of four genes of the CodY regulon was altered in *ilvD* mutants relative to D39N (Supplementary Figure 7), the pattern was inconsistent, with both genes negatively-regulated by CodY and those reported as positively-regulated^27^ showing increased expression in the *ilvD* mutants.

We also considered the possibility that loss of the iron-sulfur cluster containing protein IlvD might be advantageous in environments of high oxidative stress, such as the bloodstream and within phagocytic cells, due to a reduced risk of triggering damaging Fenton chemistry. We assessed oxidative stress resistance by exposing pneumococcus to either hydrogen peroxide (Supplementary Figure 8A) or paraquat (Supplementary Figure 8B). In both cases, we observed no significant difference in survival between D39N and Δ*ilvD* pneumococci. Whilst these findings demonstrate that oxidative stress resistance is unaltered in the absence of IlvD under the conditions tested, it remains possible that the assay fails to recapitulate the conditions relevant to bloodstream or phagocytic cell environments.

We reasoned that a further way in which loss of IlvD might benefit pneumococcus in the bloodstream, or within phagocytic cells, might be by enabling more efficient use of limited iron. Phagocytic cells export iron or sequester it, as part of a nutritional immunity program aimed at controlling bacterial growth^28^. Whilst the pneumococcus relies mainly upon manganese as a metal cofactor for proteins, it does have a suite of iron-sulfur cluster proteins, some of which are likely important during systemic infection. Both wild-type and Δ*ilvD* grew poorly in chemically defined media containing the iron chelator bathophenanthroline disulfonate, confirming a requirement for iron in pneumococcus (Supplementary Figure 9). When we assessed survival of D39N and the *ilvD* deletion strain within phagocytic cells that had been pre-loaded with iron, we recovered similar numbers of viable bacteria for each strain (Supplementary Figure 10), suggesting the fitness benefit of losing *ilvD* is only realised in iron-restricted environments. This is consistent with a model in which, in the absence of IlvD, available iron can be redirected to Fe-S cluster proteins specifically required under systemic infection conditions.

Collectively, our findings demonstrate that despite apparent BCAA auxotrophy *in vitro*, pneumococcus requires the BCAA biosynthesis machinery for efficient colonisation of nutritionally poor environments, such as the nasopharynx. In the bloodstream, where BCAAs are relatively abundant, but free iron is scarce, loss of IlvD may free up iron for incorporation into Fe-S proteins, such as the pyruvate formate lyase (Pfl)-activating enzyme or LipA, that are required for survival in harsh, anaerobic conditions.

## Discussion

Pneumococcal lineages have differing propensities to cause invasive disease, with some GPSC-serotype combinations displaying elevated attack rates^29,30^. However, large scale genomic studies have revealed few differences between bloodstream isolates and those isolated from other infection sites^31,32^. Our analysis-using matched lung and blood pneumococci from individual infections - supports this, with no fixed non-synonymous SNPs identified in the blood-borne pneumococcal population, relative to matched populations from lungs. However, we did identify low population-frequency non-synonymous mutations in blood isolates. Notable amongst these were SNPs in *ilvD*, which arose multiple times in independent populations, clustering in a short stretch of highly conserved nucleotide sequence^19^. Bacteraemia is an acute event, offering limited opportunity for advantageous mutations to sweep to fixation. Furthermore, as bloodstream infection represent an evolutionary dead-end for the pneumococcus, mutations that might prove advantageous in the systemic circulation are unlikely to be maintained in the population if they come with fitness costs in other host environments. Loss-of-function mutations in *ilvD* fit this model, conferring advantages in mouse bacteraemia models but with a substantial fitness cost in the primary niche of nasopharynx. In support of an important role for IlvD in colonisation, analysis of clinical isolates suggests the gene is under purifying selection. Nonetheless, several *ilvD* variants showed enrichment amongst clinical isolates from invasive pneumococcal disease, notably the S38P and V428I mutations, which were observed across multiple lineages. The effects of these mutations on IlvD function, and whether they contribute directly to an increased propensity to cause bloodstream infections remains unknown.

Previous reports suggest that loss of Ilv proteins can have both beneficial and deleterious consequences for microbial fitness in cell culture and mouse models, suggesting a context-specific role for BCAA biosynthesis in infection. In *S. mutans*, deletion of *ilvE*, encoding a branched-chain aminotransferase, conferred increased survival within mammalian cells, which was attributed to increased uptake of environmental BCAA and associated disruption of host cell function^21^. We did not quantify BCAA uptake in our study but did observe elevated expression of BCAA transport genes in *ilvD* mutants, which may have contributed to their increased survival within phagocytic cells. In *Staphylococcus aureus*, inactivation of BCAA uptake prohibits growth within macrophages, whilst a strain lacking *ilvD* showed upregulation of the BrnQ amino acid transporter, helping to sustain intra-macrophage viability by importing amino acids from the extracellular environment^14^. Environmental scavenging of BCAAs may be a common requirement for microbial species to survive within intracellular environments. In pneumococcus, deletion of *ilvC* has been shown to attenuate virulence in a peritonitis model and to reduce expression of both *ply* and *lytA*. We observed modestly reduced virulence in a pneumonia model but increased competitive fitness in a bacteraemia model for an *ilvD* deletion strain, suggesting the route of infection influences the contribution of BCAA biosynthesis genes to virulence. This likely reflects different BCAA bioavailability within different infection environments. BCAAs are found at concentrations of up to 800 µM in blood^33^, whilst airway concentrations are notably lower. In the mouse airways, we previously reported concentrations of 200 nM in lungs and 80 nM in nasopharynx ^4^. The requirement of pneumococcus for IlvD for prolonged nasopharyngeal carriage likely reflects the limited environmental BCAA supply within this environment.

Several Gram-positive species have been reported as BCAA auxotrophs, including pneumococcus and *S. aureus*^34^. These findings may be an artefact of *in vitro* conditions used for growth, as both species encode the entire suite of proteins required for *de novo* BCAA biosynthesis within their genomes. The activity of the *ilv* operon in Gram-positive bacteria is controlled by the conserved global regulator CodY, which acts as a repressor. Deletion of CodY can release repression and induce BCAA synthesis in *S. aureus*^14,35^. Comparable experiments in pneumococcus are complicated by the essentiality of CodY^36^, with reported *codY* deletion strains likely containing compensatory mutations elsewhere in the genome. Our finding that IlvD is required within the nasopharynx and that BCAA supplementation restored the ability of Δ*ilvD* to compete in this environment leads us to conclude that *de novo* BCAA synthesis is necessary for prolonged nasopharyngeal carriage. An unidentified *in vivo* cue may be required to derepress *ilv* genes, which has not been captured by the *in vitro* conditions used here or in previous studies that suggested BCAA auxotrophy in pneumococcus.

Our initial efforts to rationalise the fitness benefit of IlvD loss in bloodstream environments, and particularly within phagocytic cells, focused on oxidative stress responses, given the presence of a 4Fe-4S prosthetic group within the protein. Stress-induced Fe-S cluster damage can trigger the stringent response in multiple bacterial species and contributes to phagocyte-mediated bacterial killing^37–40^. Ferrous iron (Fe²⁺) can react with hydrogen peroxide via Fenton chemistry, generating hydroxyl radicals that damage DNA and impair bacterial viability^41^. We reasoned that loss of IlvD might therefore increase tolerance to oxidative stress. However, we observed no significant changes in survival of wild type versus Δ*ilvD* pneumococci following exposure to hydrogen peroxide or paraquat. Pneumococcus is highly resistant to oxidative stress, having evolved multiple regulatory mechanisms to reduce toxicity associated with hydrogen peroxide produced by the actions of its pyruvate oxidase, SpxB^42^. However, it remains possible that oxidative stress resistance might be altered in *ilvD* mutants within the intracellular environment of phagocytic cells.

Obtaining iron is a challenge for bacteria in infection contexts, as evidenced by the presence of siderophores and other iron-scavenging proteins in many pathogen genomes^43^. Iron sequestration is a major defence mechanism of nutritional immunity and phagocytic cells produce iron scavengers, including lactoferrin, or store iron in dense crystalline form in ferritin cages^44^. Iron-loading of THP-1 cells neutralised the fitness advantage shown by the Δ*ilvD* strain, as compared to wild type, within the intra-phagocyte environment. This suggested to us that the benefit of IlvD loss might relate to the freeing up of iron for other Fe-S cluster-containing proteins. The pyruvate formate lyase activating enzyme PflA, which supports anaerobic metabolism^45^, is an Fe-S protein that might be important within the oxygen-depleted environment of phagocytic cells. The Fe-S cluster-containing lipoate synthase LipA may also be necessary to maintain central metabolism under such conditions, given the links between lipoic acid metabolism and oxidative defence^46^.

This study highlights the importance of the BCAA biosynthesis machinery in *S. pneumoniae* lifestyles. Pneumococcus has retained a protein that is a liability within some host environments – most strikingly within phagocytic cells – due to the benefits it confers within the primary niche of nasopharynx. Biosynthesis of BCAAs appears to be an important contributor to fitness in the nutrient-restricted environment of the airways.

## Methods

### Bacterial strains and growth conditions

All *S. pneumoniae* used in the study were derivatives of the serotype 2 D39 strain, with mutants prepared from our in-house stock, D39N (NCBI Accession number: CP061208) (Table 2). Frozen bead stocks were first streaked on a gentamicin-blood agar base (BAB) plate and allowed to grow overnight at 37°C in a 5% CO_2_ incubator. The purity of the culture was confirmed by sensitivity to optochin. Liquid cultures were grown in brain heart infusion (BHI) broth without shaking at 37°C in a 5% CO_2_ incubator. Some broth culture experiments were performed in chemically defined media (CDM) containing varying concentrations of BCAA^47^.

**Table 2.** Strains used in or generated for this study.

| **Name** | **Source** | **Description** |
| --- | --- | --- |
| D39N | Previous study<sup>17</sup> | In-house stock of lab strain D39 |
| D39N<sup>SmR</sup> | Previous study<sup>17</sup> | *rpsL* point mutation yielding K56T amino acid substitution |
| $\Delta ilvD$ | This study | Clean deletion of *ilvD* gene |
| IlvD<sup>S273C</sup> | This study | *ilvD* point mutation yielding S273C amino acid substitution |
| IlvD<sup>T274P</sup> | This study | *ilvD* point mutation yielding T274P amino acid substitution |
| IlvD<sup>N275I</sup> | This study | *ilvD* point mutation yielding N275I amino acid substitution |
| IlvD<sup>T277N</sup> | This study | *ilvD* point mutation yielding T277N amino acid substitution |
| IlvD<sup>L278I</sup> | This study | *ilvD* point mutation yielding L278I amino acid substitution |

### Genome sequence analysis of blood isolates from a mouse experimental evolution study

Populations of pneumococci were evolved from D39N via serial passage of lung-resident bacteria in a mouse pneumonia model^17^. Sequence data associated with the lung-resident populations and from matched populations recovered from blood are accessible via NCBI Bioproject PRJNA658145 under SRA submission number SUB16215073. Mutations identified in the lung-resident population are described in an earlier publication^17^. Genomic DNA was sequenced using the Illumina platform at the Sanger Institute. Pairwise comparisons of lung and blood populations from individual mice were performed, with variant calling versus the D39N ancestor, using BreSeq in population mode^18^.

### Bacterial mutant strain construction

In total, eight strains were used in this study. In addition to D39N wild type, a streptomycin-resistant D39N strain was used, generated via introduction of a point mutation in *rpsL* (producing a K56T substitution). All remaining mutants were generated using the PhunSweet system, with positive selection on 1 μg/mL erythromycin agar and counter-selection with 15 mM chlorinated-phenylalanine, 10% sucrose. The cassette was then replaced to generate clean mutants, which were confirmed by Sanger sequencing. Oligonucleotide primers used in the construction of strains and in gene expression analysis are listed in Table 3.

**Table 3.**
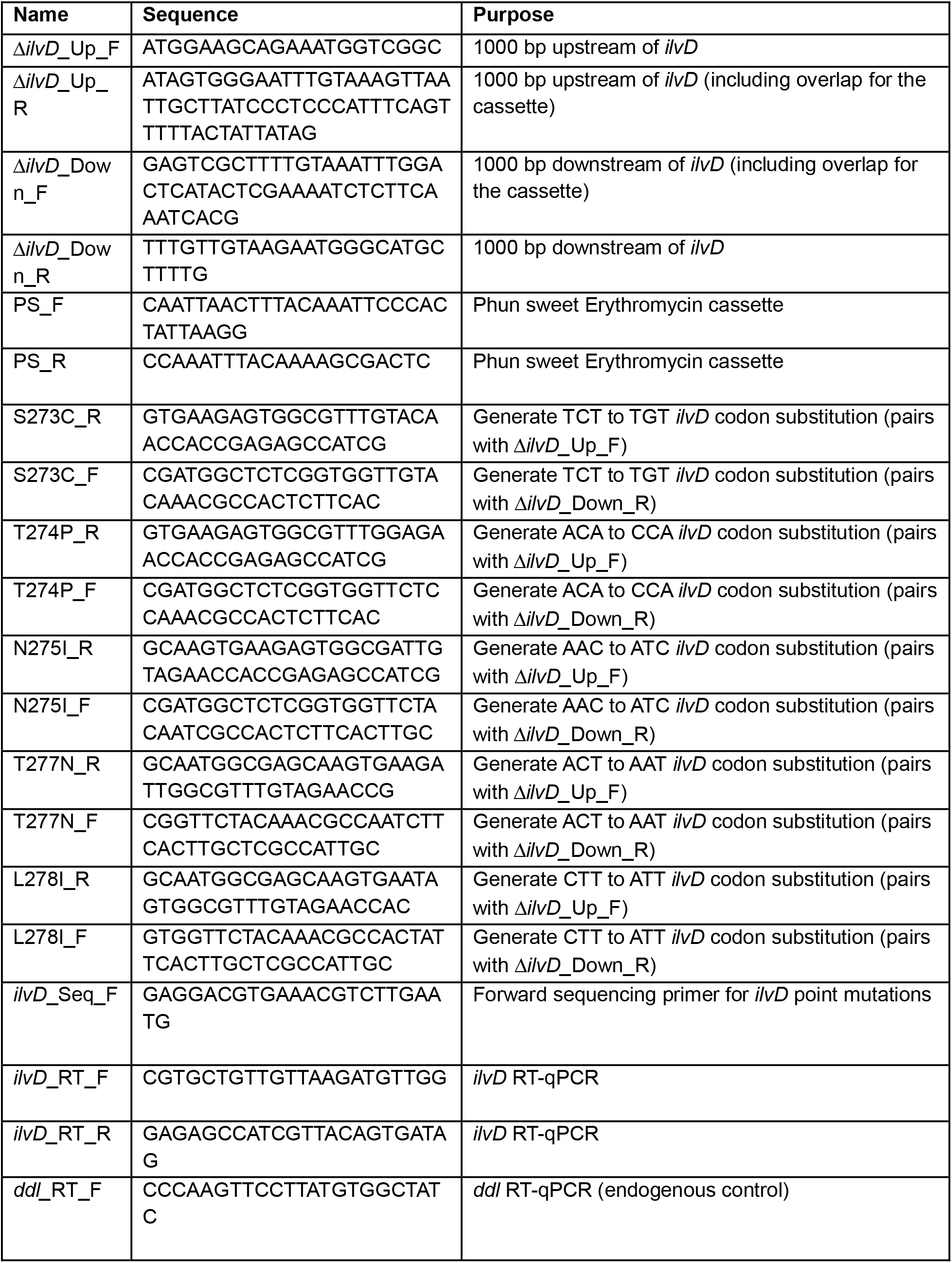
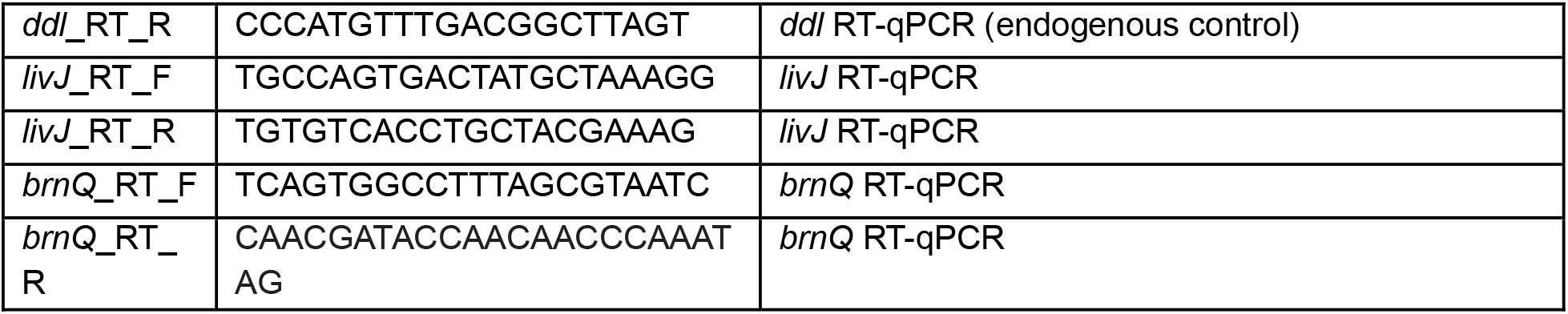
Oligonucleotide primers used in this study.

### Gene expression analysis

Primary cultures of bacterial strains were grown overnight in BHI. On the following day, cultures were pelleted, washed in phosphate buffered saline (PBS) and resuspended in either fresh whole human blood or BHI. The cultures were further incubated for 2 hours, and RNA purification was performed using the RNA-Bee/Stat60-based manual method (amsbio). DNA carry-over was digested using the Turbo DNA-free kit (Invitrogen), according to manufacturer’s instructions. The RNA concentration was quantified at OD260 using the NanoDrop8000UV–vis Spectrophotometer (Thermo Scientific). Purity was determined by 260/280 nm ratio (target 1.8-2.0). A total of 500 ng RNA was converted to cDNA using the Promega cDNA synthesis kit. The cDNA was stored at −20°C until further use. A no reverse transcriptase (NRT) control was included for assessment of DNA contamination. Experiments were performed in biological duplicates and for each replicate, a total of 3 reactions were set with the GoTaq® qPCR Master Mix (Promega), as per manufacturer’s instructions. Reactions contained 1 μl cDNA and 0.2 μM forward and reverse primers (Table 3). Template-free and DNA polymerase-free controls were included in each assay run. PCRs were performed on the Applied Biosystems GeneAmp 9700 PCR system using MicroAmp™ fast optical 96-well reaction plates (Applied Biosystems) under the following conditions: 2 min at 95°C followed by 40 cycles of 15 sec at 95°C and 1 min at 60°C. Analysis of relative gene expression used the 2^-ΔΔCt^ method with *ddI* used as the housekeeping gene for normalisation of expression. Oligonucleotide sequences for the primers used in the study are enlisted in Table 3.

### Phagocytosis assays

The fitness of pneumococcal strains associated with or inside phagocytes was tested in the THP-1 monocyte cell line (ATCC TIB-202). THP-1 cells were cultured at 37°C and 5% CO_2_ in RPMI (Gibco) supplemented with 10% fetal bovine serum (FBS) and Penicillin-Streptomycin antibiotic cocktail. For differentiation, THP-1 monocytes were treated with 50 ng/mL Phorbol 12-myristate 13-acetate (PMA, Thermofisher Scientific) for 24 hours, at a density of 1×10^5^ cells/well in a 96 well F-bottom plate, followed by resting in fresh media for another 24 hours, without PMA. One hour before the experiment, the media was again replaced with RPMI. Each well was infected with 1 × 10^7^ colony forming units (CFU) of pneumococcus, prepared from frozen stocks, to achieve a multiplicity of infection (MOI) of 100, and incubated at 37°C and 5% CO_2_. At 40, 60 or 120 minutes post-infection, planktonic and loosely adherent bacteria were removed using three PBS washes, and then macrophages were lysed using 0.1% triton. Serial dilutions of lysates were made in PBS and spotted on BAB plates. On the following day, the colonies were enumerated by CFU count.

For the iron-supplemented phagocytosis assay, RPMI was supplemented with 5 µM iron. In these experiments, the bacterial infection stocks were adjusted to an MOI of 100 in the same iron-supplemented RPMI, and the cells were infected for 2 hours, as previously described. Cells were lysed using Triton X-100 at different time points, and bacterial survivors were enumerated by plating on blood agar plates.

### Assessment of intracellular survival by flow cytometry with pHrodo dye

The fitness of mutants inside THP-1 cells was assessed using flow cytometry. The cell seeding and differentiation protocol were as described for phagocytosis assays. Prior to THP-1 infection, frozen stocks of bacterial strains were incubated with 1 mM pHrodo dye for 90 minutes in the dark at room temperature. Following staining, bacterial numbers were adjusted to 1 × 10 CFU (MOI of 10) and used to infect THP-1 cells for 60 minutes. Unstained bacteria controls were included in all assay runs. The phagocytosis-inhibitor cytochalasin D at 1 µM concentration was included in some wells as an additional control. After 60 minutes of incubation, the extracellular bacteria-containing media was removed, and 50 μL of Gibco TrypLE was added to each well to detach THP-1 cells. Plates were then incubated at 37°C with 5% CO₂ for 30 minutes. Subsequently, 150 μL of PBS was added and mixed vigorously to ensure complete detachment of the cells. Finally, the samples were run through a Novocyte flow cytometer, quantifying pHrodo fluorescence in the R675-A channel. Data analysis was undertaken using FlowJo software.

### Growth competition assays

All tested strains were streaked on BAB plates from frozen stocks and grown overnight at 37°C in 5% CO_2_. The following day, 2-3 colonies from each plate were used to inoculate 2 mL of BHI broth or CDM, which was incubated overnight at 37°C in 5% CO_2_. Before setting a competition assay, the OD_600_ of the strains was adjusted to 0.05. Mutants were competed individually against D39N^SmR^ by co-culture in different environmental conditions, namely human serum, BHI and BCAA-restricted CDM (containing 5 µM BCAA, a 500-fold reduction from standard CDM conditions). 50 μL of each bacterial culture (Sm^R^ WT and the test strain) was transferred to 1.4 mL human serum/BHI/CDM-containing tubes. The cultures were further incubated at 37°C for 8 hours in 5% CO_2._ The time was increased to 24 hours in BCAA-deficient CDM to account for slower growth in BCAA-restricted conditions. After incubation, serial dilutions were made in PBS and plated onto BAB and streptomycin (800 μg/mL)-containing BAB plates to differentiate between D39N^SmR^ and mutant colonies. The number of colonies were counted, and the CFU for each strain was calculated. CFU counts for D39N^SmR^ were calculated from the number of colonies on the streptomycin plates. CFU counts for the test strains were calculated from the total number of colonies on the BAB plates minus the number obtained from the streptomycin plates. Before testing mutant strains in competition assays, the D39N wild type was first competed against D39N^SmR^ under the different environment conditions, to account for any fitness cost associated with the *rpsL* mutation. Competition indices (CI) for subsequent competitions between mutants and D39N^SmR^ were thereafter adjusted to account for this fitness coefficient.

### *In vitro* growth curves

A sweep of colonies from a BAB plate was inoculated into 2 mL of BHI broth and grown overnight at 37°C in 5% CO_2_. On the following day, the OD_600_ was diluted to 0.002, and 200 μL of the bacterial culture was added to a 96-well microplate, with at least 3 replicates per strain. BHI broth was used as a blank control. Plates were incubated for 18 or 24 hours, and readings were taken every 15 minutes at OD_600_ on a Tecan plate reader. The iron chelation growth curves were performed in chemically defined media made without any extra iron supplementation, hence milli Q water was the only iron source. The iron-deficient conditions were induced using the Bathophenanthrolinedisulfonic acid chelator at 300 µmol/L. The cultures were adjusted to 0.002 OD_600_ and growth was assessed by taking reading every 15 minutes in a Tecan plate reader.

### Capsule thickness determination

Capsule thickness was determined by a FITC-dextran exclusion assay adapted from Eichner *et al*., 2025^48^. Staining solution (2mg/mL FITC-dextran and 10 µM Nile red in PBS) and agarose-dye mix (2mg/mL FITC-dextran and 10 µM Nile red in 1% ultra-pure agar, kept liquid at 50 ^◦^C) was prepared on day of imaging. 100 µl of agarose-dye mix was pipetted into the centre of a gene frame and a second slide used to make an agar pad. Strains were cultured in 10mL BHI overnight to stationary phase. 500 µL of culture was pelleted by centrifugation at 3000 xG for 3-minutes at room temperature and washed three times in 1mL PBS. During the second wash 1 µL DAPI was added and incubated rolling at room temperature for 5-minutes. Cells were pelleted and resuspended in 100 µl of staining solution. 7-10 µl of resuspended cells was pipetted on to the agar pad and cover slip applied. Samples were imaged for up to 3-hours post-preparation. Images were acquired on a Leica Stellaris 8 confocal inverted microscope. Images were analysed in Fiji (ImageJ version 2.14.0/1.54f) using a custom macro detailed in Eichner *et al*., 2025^48^.

### Oxidative stress resistance assays

For hydrogen peroxide sensitivity assay, the mid-log phase pneumococci were added in duplicates to a flat-bottom microtiter plate wells and incubated at 37 °C for 30 min in BHI with 25, 37.5 and 50 mM of hydrogen peroxide before making dilutions for plating. The CFU/mL of each isolate was determined via Miles and Misra dilution onto gentamicin BAB plates and the percentage killing was calculated for each test isolate in comparison to the untreated (BHI only) control.

For paraquat sensitivity assay, 1 mL of overnight cultures were spun at high speed and adjusted to 0.01 OD₆₀₀. Following that, 140 µL of the normalised cultures were added to a flat-bottom 96-well plates. Paraquat-treated wells received 60 µL paraquat solution (200 mM stock, final concentration 60 mM), while control wells received 60 µL sterile water. For survival determination, 20 µL from each well was sampled and serially diluted in the PBS before spotting on blood agar plates. Colonies were counted the following day, after overnight incubation and CFU/mL was calculated.

### Mouse and Drosophila infection studies

Animal experiments were performed with prior approval of the UK Home Office and the relevant Animal Welfare and Ethical Review Boards. Experiments were performed under project licenses PP2072053 and PP4318217, at the University of Dundee. The principles of the Declaration of Helsinki were observed throughout. Mice were housed in individually ventilated cages, with access to food and water *ad libitum*. Environmental enrichment was provided in all cages and mice were acclimatised to the animal unit for 9-10 days before use. Mice were randomly allocated to cages on arrival in the animal unit by staff with no role in study design. For experiments reported in this manuscript, individual mice were considered as the experimental unit. Sample sizes, controls and statistical analyses are detailed in the figures and accompanying legends.

CD1 outbred female mice between the ages of 6–8 weeks (Charles River, Oxford, UK) were used in the study. To prepare a standard inoculum, pneumococci were grown to OD_600_ ∼1 in BHI supplemented with 20% FBS. The infection stocks were kept frozen until needed. For intravenous route time-point infections (bacteraemia model), the CD1 mice were kept in a restraining apparatus and 50 μL containing 1 × 10^6^ CFU resuspended in PBS for each strain was injected through a tail vein. Mice were tail bled at different time points (8, 12 and 24 hours) and euthanised after 24 hours to harvest blood and organs for bacterial titre quantification. Organs were processed using a hand-held tissue homogeniser and dilutions were made in PBS before spotting on BAB plates. Colonies were counted the following day. For bloodstream competition experiments, mice were intravenously administered with a single dose containing a 50:50 mix (5×10^5^ CFU each) of D39N^SmR^ and D39NΔ*ilvD*.

For the pneumonia model, mice were intranasally infected under light anaesthesia, using a mix of oxygen and isoflurane (2.5%), with 1 × 10^6^ CFU of each strain in 50 μL PBS. At different time points (8, 12 and 24 hours), blood was collected from tail veins for bacterial titre quantification. At 24 hours, all mice were euthanised, and organs were collected for CFU comparison.

Nasopharynx competitions were performed in a mouse colonisation model in which a 10 μL volume of PBS, containing 5×10^4^ CFU each of D39N^SmR^ and D39NΔ*ilvD,* was intranasally administered to mice under light anaesthesia. The mice from each cage were humanely sacrificed at pre-determined time points post infection, as detailed in figure legends. The bacterial load was measured in the excised nasopharynx. In all competition experiment, the CFU counts for the D39NΔ*ilvD* strain were calculated by subtracting the colony counts of tissue plated on streptomycin BAB plates from the total CFU counts on non-selective BAB plates.

Drosophila infections were carried out as has been described previously^24^. Briefly, *S. pneumoniae* were grown to mix-exponential phase in BHI. Bacteria were then pelleted and resuspended in PBS at OD600 = 0.01. 50 nl of this suspension was injected into the hemolymph of 5-10 day old male *w^1118^ Drosophila melanogaster.* After infection, flies were kept at 29°C on media containing 10% autolysed brewer’s yeast, 8% fructose, 0.8% agar, supplemented with propionic acid and nipagin.

### Histology analysis of pneumococcal plaques

Analysis of brightfield H&E slide scan images from pneumococcus-infected and uninfected mice was performed using QuPath version 0.5.1. Tissue detection for heart and lung was carried out using Pixel Classifiers: briefly, representative regions ∼1 mm^2^ for heart (18 regions in total) and lung (19 regions in total) were used to train classifiers in 3 rounds, the second and third correcting for errors in the previous round. The same strategy was used to train classifiers to detect plaques in heart and lung tissue (Heart: 4 rounds, 19 regions in total; Lung: 2 rounds, 17 regions in total). All final classifiers used the RTrees method. The classifier to detect heart tissue (minimum size 1mm^2^) was at low resolution (2.75 µm, scales 1,4,8) using Gaussian and Weighted deviation features. The classifier to detect lung tissue (minimum size 1mm^2^) was at very low resolution (5.5 µm, scales 0.5,1,4,8) using Gaussian and Weighted deviation features. The classifiers to detect heart and lung plaques (minimum size 0.25 µm^2^) were at very high resolution (0.34 µm, scales 1,2,4,8) using Gaussian, Weighted deviation and Hessian determinant features. Batch analysis was carried out using groovy scripts to first generate tissue region Annotations and then plaque Detections within these for all heart and lung images.

### Genomic analysis of clinical isolates

We detected absence of *ilvD*, disruptive mutations, and non-synonymous mutations in *ilvD* using ARIBA version 2.14.6 and the *ilvD* reference sequence from the D39V strain (NCBI accession number CP027540). The output is summarised and visualized using R version 2026.04.0+526 (https://github.com/StephanieWLo/ilvD). Absence and disruptive mutations from ARIBA were further confirmed by mapping the raw reads of the sample onto the ilvD reference sequence and the results were examined manually for confirmation. Absence and disruptive mutations from ARIBA were further confirmed by mapping the raw reads of the sample onto the *ilvD* reference sequence and the results were examined manually for confirmation.

The dN/dS analysis of ilvD was performed using a Bayesian method based on GenomegaMap (https://github.com/bacpop/TOMBOMBADIL_jax). The dN/dS was reported as a median with 90% credible interval for each amino acid position across IlvD. Metadata for the 24,455 Global Pneumococcal Sequencing project (https://www.pneumogen.net/gps/) genomes used for analysis in this study can be found in Supplementary Dataset 3.

## Statistical Analysis

GraphPad Prism version 10.2.0 was used for statistical analysis. Statistical tests undertaken for individual experiments are detailed in the respective figure legends. P < 0.05 was considered to be statistically significant, after adjustment for multiple comparisons. Data are presented as mean +/-standard deviation, unless otherwise stated.

## Acknowledgements

SK was supported by a Discovery Medicine North PhD studentship, funded jointly by the UK Medical Research Council (MRC), the University of Liverpool and the University of Dundee. KN is supported by a Wellcome Trust PhD studentship. DRN acknowledges project grant funding from MRC (MR/X009130/1). The authors gratefully acknowledge the University of Dundee Imaging Facility, for support with tissue histology, Fred Simeons, at the University of Dundee Drug Discovery Unit for training support and Luisa Hiller, at Carnegie Mellon University for useful scientific discussions.

## Author Contributions

S.K. and D.N. conceived and designed the study. S.K., A.G. A.W., N.M, K.N. and E.K. performed experiments. S.K., L.L., K.N., S.L. and D.N. analysed data. D.N., M.D. and S.L. provided resources and reagents and supervised staff. S.K. and D.N. wrote the manuscript, with input from all authors.

