## Supplementary Figures and Tables for "Niche-specific contribution of the branched-chain amino acid biosynthesis protein IlvD in *Streptococcus pneumoniae* infection"

Kumar *et al.*

**Supplementary Figures and Tables**

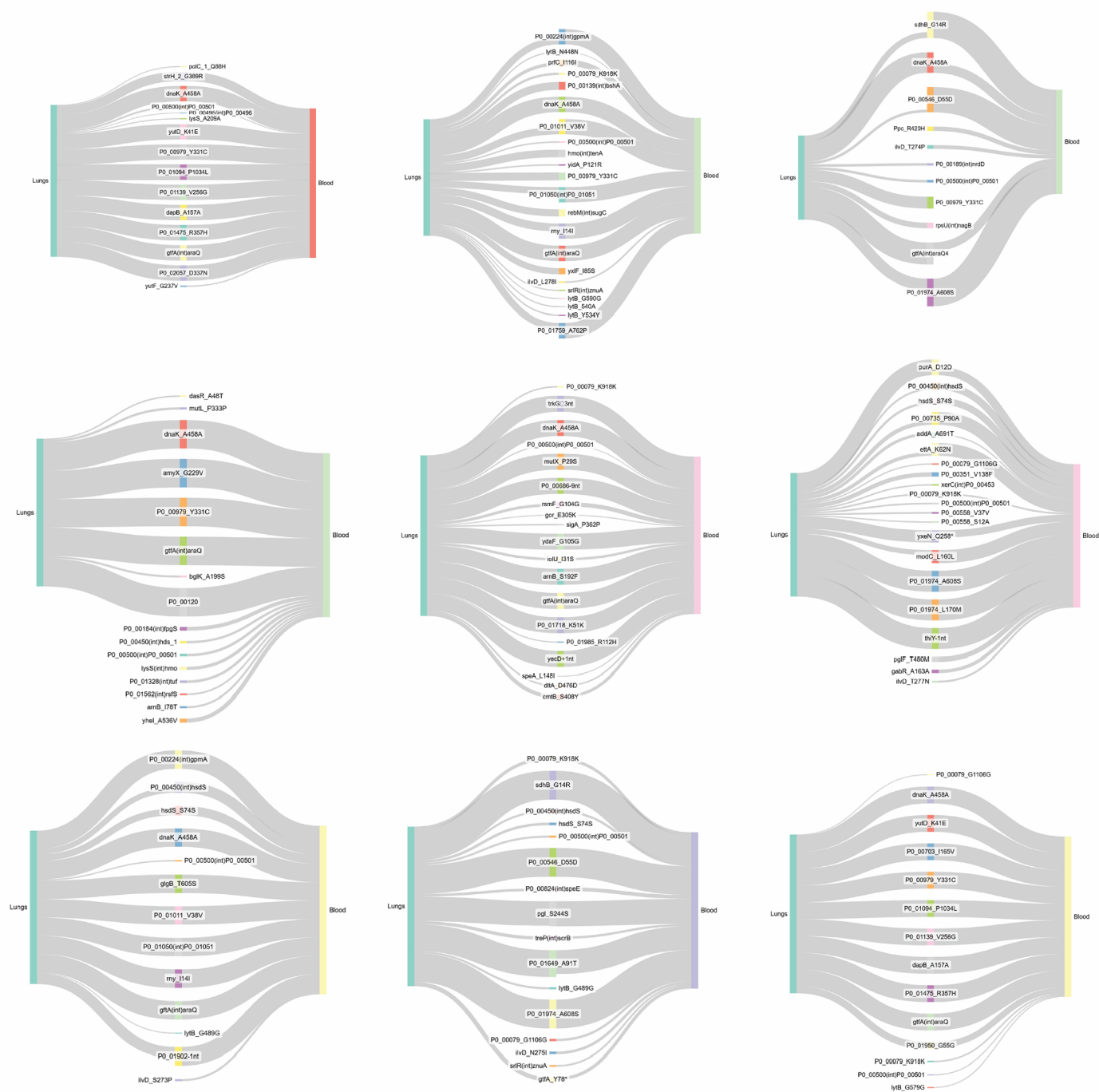

**Supplementary Figure 1. Loss and gain of mutations during the lung-blood transition in an invasive pneumococcal pneumonia mouse model.** Sankey plots showing the fate of pneumococcal genotypes identified in lungs during the progression to bacteraemia. Lines originating in lungs and terminating at the midpoint represent diversity not carried over to blood. Lines arising at the midpoint represent diversity present in blood but not identified in lungs. The thickness of the line represents the relative frequency of each mutation. The plots shown are representative example of matched lung and blood bacterial populations from nine of ten experimental-evolution lineages. An example from the tenth lineage is shown in main Figure 1B. Plots were generated using SankeyMATIC.

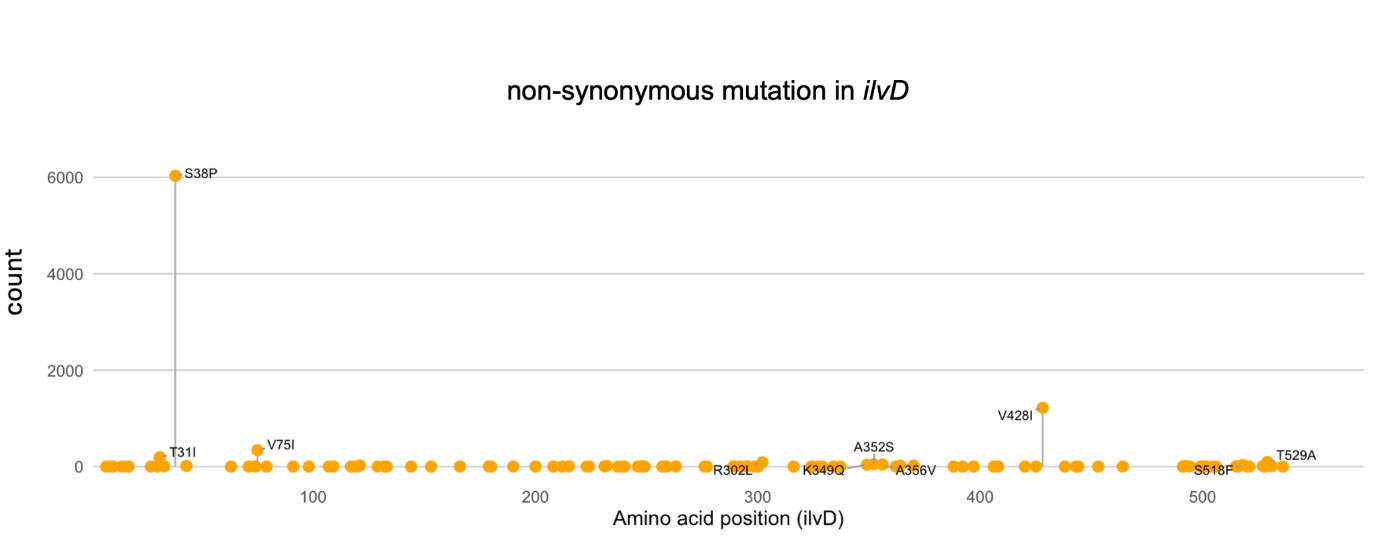

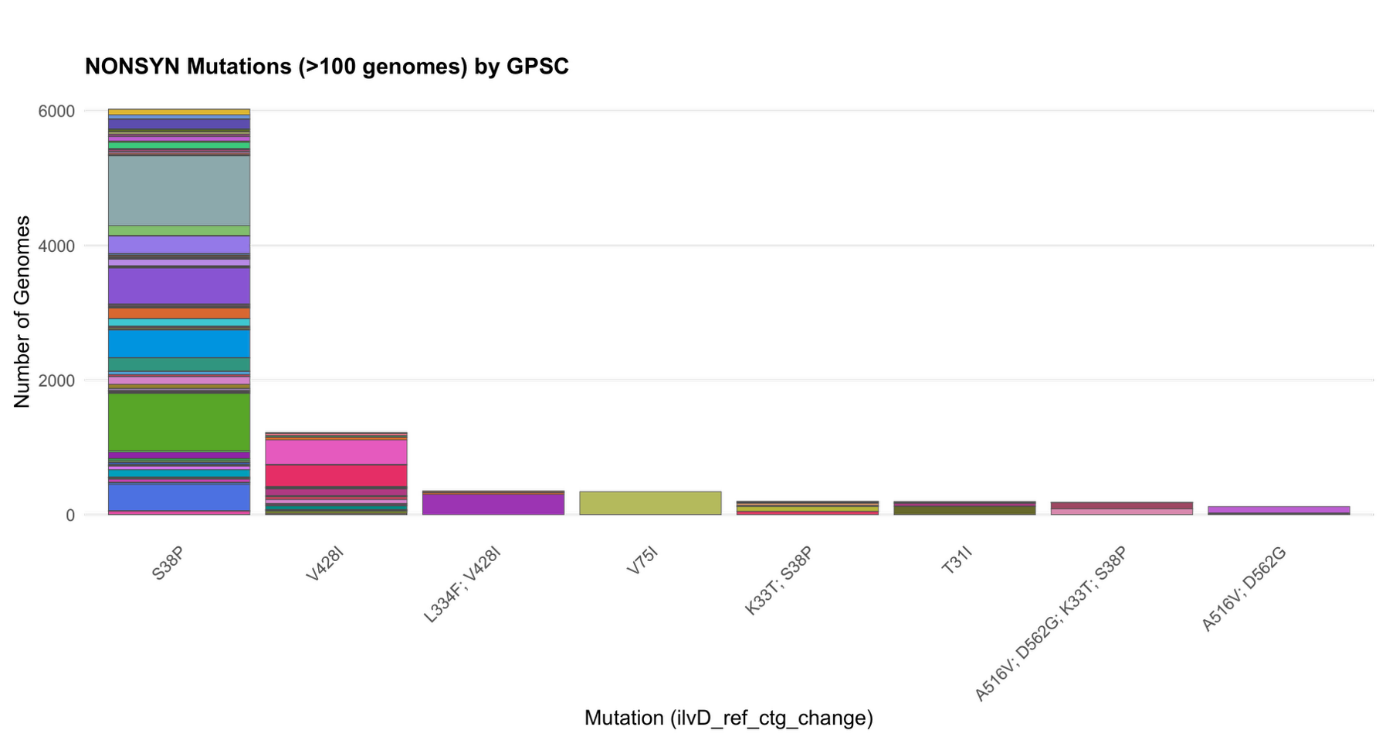

**Supplementary Figure 2. Non-synonymous mutations in *ilvD* are found throughout the coding sequence and are distributed across global pneumococcal sequencing clusters.** The top panel shows the amino acid positions of identified non-synonymous mutations in *ilvD* amongst clinical isolates, relative to the *ilvD* reference sequence from D39V (NCBI accession number CP027540). The lower panel shows mutations represented by at least 100 genomes. Banding colours represent GPSC. The V75I mutation is associated with GPSC26.

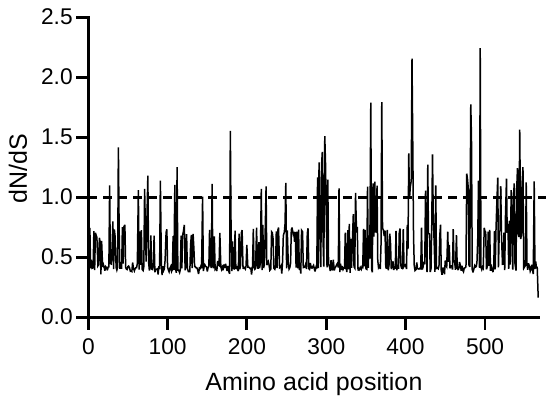

**Supplementary Figure 3. Analysis of evolutionary pressures on IlvD.** The ratio of non-synonymous to synonymous mutations (dN/dS) at each amino acid position of IlvD was determined across *S. pneumoniae* genomes from the Global Pneumococcal Sequencing project.

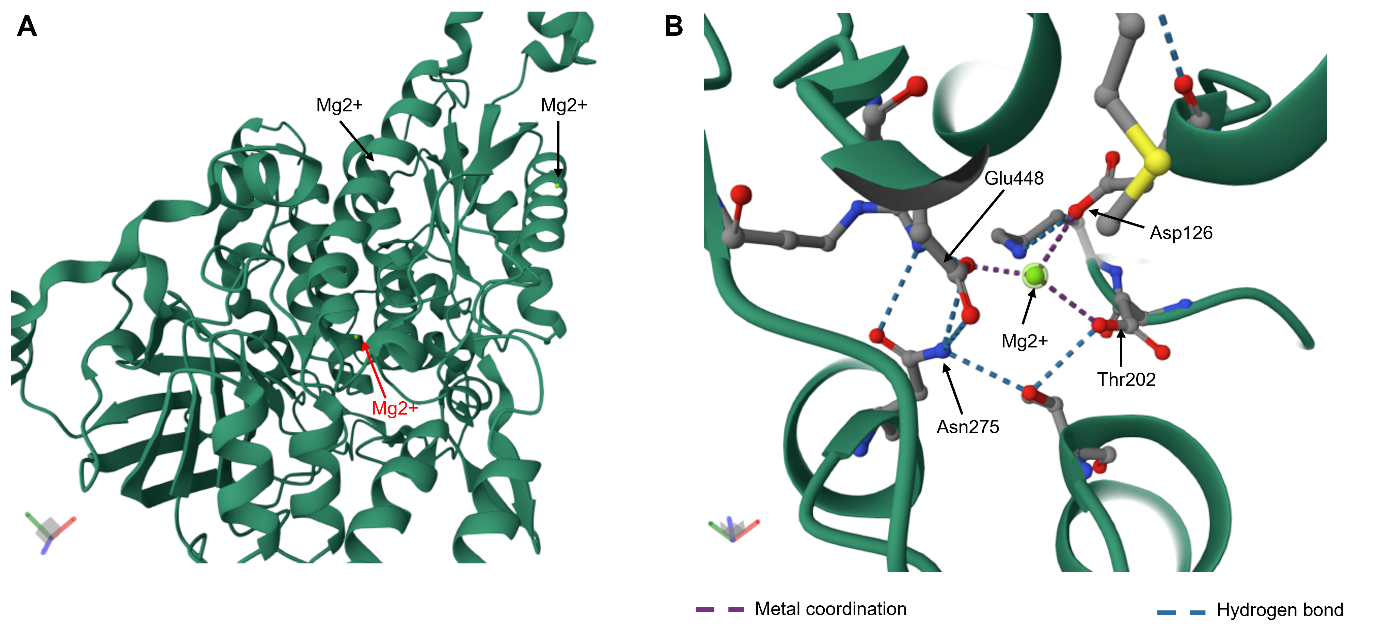

**Supplementary Figure 4. Predicted disruption of metal cofactor binding sites in *ilvD* mutants.** **A.** Predicted magnesium binding sites in *S. pneumoniae* IlvD, with position of interest highlighted in red. **B.** Interactions of magnesium with IlvD are shown, alongside stabilising hydrogen bonds at key positions, including Asn275, which was affected by a SNP in experimentally-evolved pneumococci isolated from blood. Predictions were made with alphafill, using alphafold models of Uniprot protein Q04I44.

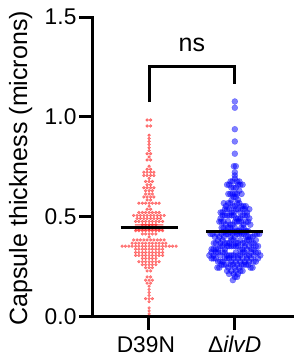

**Supplementary Figure 5. Capsule thickness is unaltered by loss of *ilvD*.** Capsule thickness was determined by FITC-Dextran exclusion assay. Each data point is an individual cell measurement and data are a composite of two independent experiments. Ns = non-significant by unpaired t-test comparison.

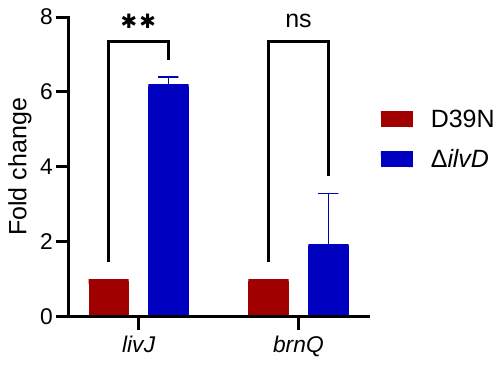

**Supplementary Figure 6. BCAA transporter expression is upregulated in *ilvD*-deletion pneumococci during culture in whole human blood.** Data are from two independent experiments, each containing four technical replicates per gene, per strain. ** = p < 0.01, ns = non-significant in two-way ANOVA with Sidak’s multiple comparison test.

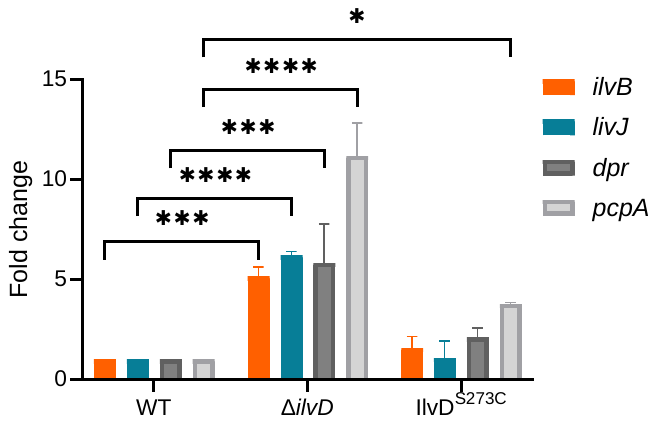

**Supplementary Figure 7. Upregulation of genes of the CodY regulon in *ilvD* mutants.** Expression of two genes reported as positively-regulated by CodY and two reported as negatively-regulated was assessed by RT-qPCR of RNA extracted from pneumococci grown in whole human blood. Data for individual genes are relative to D39N expression levels and data are from two independent experiments, each containing four technical replicates per gene, per strain. * = p < 0.05, *** = p < 0.001 and **** = p < 0.0001 in two-way ANOVA analysis with Tukey’s multiple comparison test. All remaining differences were non-significant.

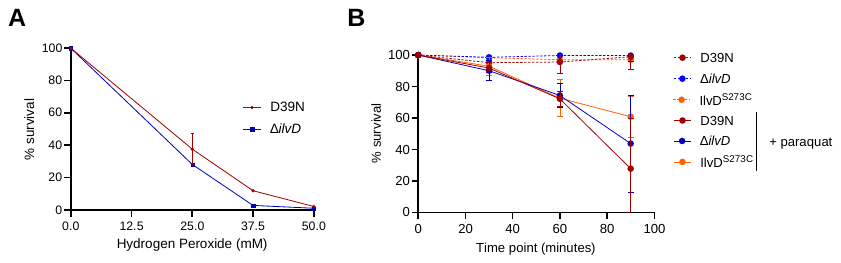
**Supplementary Figure 8. Oxidative stress resistance is unaltered in pneumococci lacking *ilvD*.** Survival of pneumococci following exposure to **A.** hydrogen peroxide at 25, 37.5 and 50 mM concentrations for 30 minutes or **B.** paraquat at 60mM concentration, assessed at 30, 60 and 90 minutes by CFU/mL. Percent survival is expressed relative to time zero. Error bars represent standard deviation of 3 biological replicates per condition.

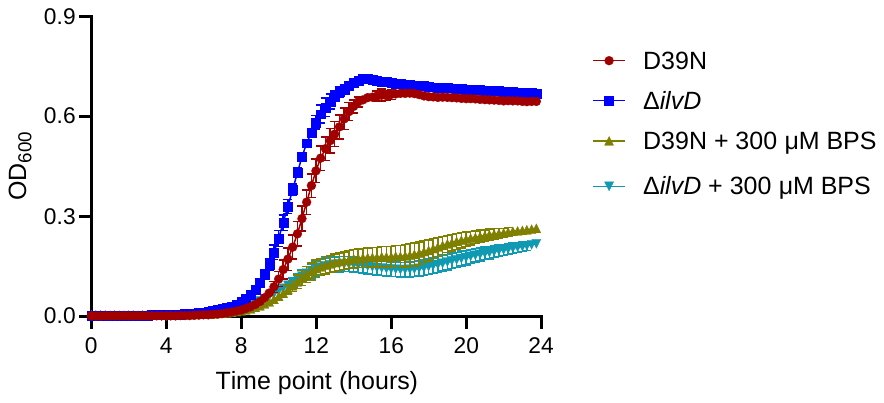

**Supplementary Figure 9. Iron is required for robust pneumococcal growth.** Strains were grown in CDM or CDM lacking iron and supplemented with iron chelator BPS. Growth was monitored over 24 hours in a plate reader. Two independent experiments were performed and data from a single representative experiment are shown.

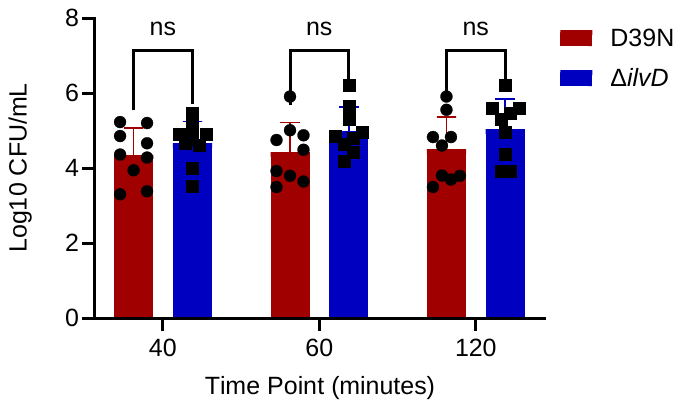

**Supplementary Figure 10. The fitness advantage of loss of IlvD within macrophages is lost when iron is not limiting.** Phagocytosis assays were performed with PMA-activated THP-1 cells cultured in iron-rich media. Surface-associated and intracellular bacterial numbers were determined at 40, 60 and 120 minutes post-infection. Data are a composite of three independent experiments, each with three technical replicates per strain, per time point. Ns = non-significant in two-way ANOVA with Sidak’s multiple comparison test.

**Supplementary Table 1. Disruptive mutations identified in *ilvD* amongst 24,455 clinical isolates of *S. pneumoniae.***

| **Isolate** | **Country** | **Source** | **Mutation type** | **Mutation*** |
| --- | --- | --- | --- | --- |
| GPS_GM_1517 | The Gambia | Nasal swab | Frameshift | K410fs |
| GPS_GM_1539 | The Gambia | Nasal swab | Nonsense | E261trunc |
| GPS_US_PATH617 | Mozambique | Blood | Nonsense | G471trunc |
| GPS_ZA_1560 | South Africa | Blood | Frameshift | G443fs |
| GPS_ZA_5681 | South Africa | Blood | Nonsense | E528trunc |
| GPS_ZA_CARRIAGE_SP458 | South Africa | Nasal swab | Frameshift | P218fs |

*fs = frameshift, trunc = truncation
